# Revisiting pangenome openness with *k*-mers

**DOI:** 10.1101/2022.11.15.516472

**Authors:** Luca Parmigiani, Roland Wittler, Jens Stoye

## Abstract

Pangenomics is the study of related genomes collectively, usually from the same species or closely related taxa. Originally, pangenomes were defined for bacterial species. After the concept was extended to eukaryotic genomes, two definitions of pangenome evolved in parallel: the gene-based approach, which defines the pangenome as the union of all genes, and the sequence-based approach, which defines the pangenome as the set of all nonredundant genomic sequences.

Estimating the total size of the pangenome for a given species has been subject of study since the very first mention of pangenomes. Traditionally, this is performed by predicting the ratio at which new genes are discovered, referred to as the *openness* of the species.

Here, we abstract each genome as a set of items, which is entirely agnostic of the two approaches (gene-based, sequence-based). Genes are a viable option for items, but also other possibilities are feasible, e.g., genome sequence substrings of fixed length *k* (*k*-mers).

In the present study, we investigate the use of *k*-mers to estimate the openness as an alternative to genes, and compare the results. An efficient implementation is also provided.

## Introduction

With the advent of high throughput sequencing technologies, the number of genomes that we have at our disposal has been steadily increasing. When a reference genome is available, sequencing and genome analysis usually involve comparing the nucleotide sequences to that reference. However, a single reference sequence can hardly account for all the variability present in nature and can not truly represent a whole species (The Computational Pan-Genomics Consortium, 2018).

In 2005 the term *pangenome* was used by Tettelin et al. (Tettelin, Masignani, et al., 2005) to describe the set of all distinct genes present in a species: either present in all genomes, defined as *core genes*, or present in just some, called *dispensable genes*. One goal was to predict how many additional genomes should be sequenced to fully describe a bacterial species. While the concept of pangenomes was later extended beyond bacteria, to plants and animals, one of the most outstanding discoveries at the time was that some species possess an *open* pangenome and others a *closed* pangenome.

For an open pangenome, the predicted number of genomes that need to be sequenced in order to get a full picture of the species is large or even undetermined. New distinct genes are found each time a newly sequenced genome is added. A closed pangenome, instead, needs much fewer genomes to portray the species. A binary classification of pangenomes into open and closed categories often proves to be challenging and multiple studies have argued against such a definitive separation (Gautreau et al., 2020; Cummins et al., 2022; Tonkin-Hill et al., 2023). We refer to *openness* to describe this variability and resort to use the terms open or closed only for historical reasons or comparison with other works.

In this paper, we define a genome as a set of abstract items. Possible choices of items include, but are not limited to, genes, ORFs or genome intervals of fixed size. The pangenome is defined as the union of these sets. The estimation of the pangenome *openness* requires the computation of the pangenome growth (Tettelin, Riley, et al., 2008).

The pangenome growth is computed starting with one genome and sequentially performing the union with new genomes from the set until all genomes have been considered. It follows that the final total size of the pangenome is independent of the ordering in which we choose the genomes, but the pangenome growth is not. To make it independent of the order, we can compute the pangenome growth for every possible order of the genomes, and then calculate an average pangenome growth. This procedure can be simplified by computing the average over all possible combinations of *m* genomes, where *m* goes from one to the total number of genomes. While it is possible to compute the average pangenome growth (hereafter referred to simply as *pangenome growth*) for a small number of genomes – e.g., with the eight isolates of *Streptococcus agalactiae* used in (Tettelin, Masignani, et al., 2005) – it is clear that the computation cannot scale with bigger datasets. Even if a more complex solution has been proposed to minimize the number of combinations to consider (Zhao, Jia, et al., 2014), the most common and practical solution is to randomly sample multiple permutations of the ordering of genomes.

In this work, we investigate the use of *k*-mers, short genome intervals of constant length *k*, as items for determining pangenome openness and compare to the gene-based approach. Expressing genomic sequence content through *k*-mers is a well-established approach and examples of their use can be found in many different applications, like genome assembly (Compeau et al., 2011), read mapping (Xin et al., 2013) and metagenomics (Wood and Salzberg, 2014). One of the advantages of using *k*-mers is that they require only the genome sequence, avoiding several potentially expensive and erroneous preprocessing steps needed by the gene-based approaches. For example, genome assemblies must be available or computed when using a gene-based approach, while *k*-mers can be extracted directly from sequencing reads. Moreover, in the absence of genome annotation, each sample must be annotated in silico. Lastly, the comparison between gene sequences requires an adequate definition and computation of homology, and, based on that, a definition and computation of gene families. In summary, the lack of preprocessing steps, coupled with the existence of fast algorithms for handling *k*-mers (Cheng et al., 2021; Kokot et al., 2017; Marçais and Kingsford, 2011), make the estimation of the openness based on *k*-mers fast and straightforward.

The remainder of this paper is organized as follows. Section “Related work” motivates our choice of the gene-based methods for the estimation of the openness for comparison, detailing the clustering methods employed by the tools: Roary (Page et al., 2015), Pantools (Sheikhizadeh et al., 2016) and BPGA (Chaudhari et al., 2016). Section “Methods” defines the pangenome growth function, gives an efficient method for its computation, and presents our implementation, Pangrowth. Section “Results” contains the empirical results of the *k*-mer-based and three gene-based approaches on twelve bacterial pangenomes to compare their predicted openness. Section “Conclusion” concludes the text.

## Related work

A number of approaches have been published to compute pangenome growth. Vernikos (2020) provides a useful starting point, enumerating an extensive list of tools and highlighting five notable ones: BPGA (Chaudhari et al., 2016), Roary (Page et al., 2015), LS-BSR (Sahl et al., 2014), PanOCT (Fouts et al., 2012), and PGAP (Zhao, Wu, et al., 2012). From this selection, we incorporated BPGA and Roary into our analysis. LS-BSR and PanOCT were excluded due to their considerably slow running times (see running times comparison in Page et al. (2015)). PGAP, despite its relevance in the analysis of microbial genomes, was also excluded as it does not directly support the openness analysis.

Among the remaining tools, we decided to not use PANINI (Abudahab et al., 2019), a web-based application for pangenome analysis that relies on Roary’s output, to avoid redundancy. Similarly, PanGP (Zhao, Jia, et al., 2014), despite offering an improved method to calculate the average of the permutation, was not included as it requires the pan-matrix – a binary matrix with presence absence of genes in each genome – as input. PanX (Ding et al., 2018), while being designed for pangenome analysis, lacks of explicit support for pangenome openness. PGAT (Brittnacher et al., 2011), a web-based tool for the analysis of microbial genomes, even if it supports database queries to identify genes that are present or absent in the pangenome, also does not directly report the openness. Moreover, PanACEA (Clarke et al., 2018), a pangenome tool to explore chromosome and plasmids, was excluded for the same reason. On the other hand, we decided to include in our study Pantools (Sheikhizadeh et al., 2016) since it directly supports the computation of the openness.

Therefore, hereafter we describe the clustering methods used in Roary, Pantools and BPGA in more detail, in particular how they cluster genes into gene families, which is the main difference between gene-based methods for pangenome growth estimation.

### Roary

The gene clustering approach chosen in Roary consists of four steps. First, coding sequences translated from each genome are clustered using CD-HIT (Fu et al., 2012). In this step, CD-HIT is applied iteratively with decreasing similarity thresholds. At each iteration, every cluster of genes that contains as many genes as genomes is filtered out. The clustering process of CD-HIT goes as follow. First, sequences are decreasingly sorted based on their length. The first sequence is defined to be the representative of the first gene cluster. Each sequence is then compared to all the previous gene representatives. If it is similar enough to one of these genes, it is added to the cluster, otherwise a new gene cluster is created with this sequence as the representative. Similarities between sequences are calculated using local alignment on amino acid sequences. All the proteins that are not filtered out after the last CD-HIT iteration are further processed in the second and third step. In these two steps, an all versus all comparison is performed using BLASTP, and the calculated scores are used to cluster the proteins using MCL (Enright et al., 2002). The final step of Roary is to post process the clusters, for example distributing paralogs into separate groups.

### Pantools

Similar to Roary, Pantools clusters proteins by computing their local alignments and then running MCL. To reduce the number of comparisons, an alignment is explicitly computed only when the number of shared hexamers (substrings of length six) of the two proteins can guarantee a minimum alignment score. If the similarity score of the local alignment, normalized to be independent of the length, is greater than a second threshold, an edge between the two proteins is added to a similarity graph. For each connected component in the resulting graph, a similarity matrix is computed. Each score is further scaled based on the species of origin of the considered proteins. MCL is then applied to each similarity matrix.

### BPGA

BPGA provides three options for clustering: USEARCH (Edgar, 2010) (which is the default), CD-HIT (Fu et al., 2012) or OrthoMCL (Li et al., 2003). Specifically, USEARCH uses the UCLUST algorithm (Edgar, 2010) to group genes into clusters. Within BPGA, the cluster_smallmem command is executed, a memory-efficient variant of UCLUST. The algorithm focuses on identifying centroids, which are sequences selected to represent a gene family. Other sequences that align with a centroid at or above a predefined threshold are assigned to that centroid’s family. In BPGA, genes must be converted into amino acids before clustering and are subsequently sorted by decreasing length. This ensures that the longest sequences are likely to be chosen as centroids. As UCLUST processes the genes, it constructs a database of centroids. Sequences within this database are prioritized based on the number of unique substrings of fixed length they share with the query sequence. This increases the probability of obtaining a strong match early in the list of sequences. The algorithm terminates after a fixed number of rejections or accepted sequences. The similarity between sequences is computed by aligning each string with a seed extension technique and banded dynamic programming to improve performance.

## Methods

In this section, we abstract the notion of genome to a set of *items*. Items have to be of the form that genomes may share items, while items can also be unique to genomes. Genomes are described as a subset of the universal set of items *𝒦* present in a species, or other taxonomic rank. The set *𝒦* contains the union of all the items appearing in the pangenome plus all the unseen items that may appear in some genomes not included yet. The problem of estimating the size of *𝒦* can be reduced to a well studied problem in information theory, where the increment in the total number of items is studied as a function of the number of *objects* (here, genomes) contemplated. This function is known to follow a power law, referred to as Heaps’ law (Heaps, 1978). The exponent of the power law defines the openness of the species (Tettelin, Riley, et al., 2008).

### Definitions

Let *𝒢* = {*G*_1_, …, *G*_*n*_} be a set of genomes, *P*(*G*) the power set representing all possible subsets of *𝒢*, and *𝒢*_*m*_ = {*𝒢*^*′*^ ∈ *P*(*G*) | | *𝒢*^*′*^| = *m*} the set of subsets of *𝒢* of cardinality *m*. Let *M* be a binary matrix possessing a column for each genome and a row for each item. If item *x* is present in genome *i* then *M*_*x,i*_ = 1, otherwise *M*_*x,i*_ = 0. *M* is usually referred to as *pan-matrix*.

In order to study the openness of a species, we want to examine the size of the union of genomes when considering more and more of the species’ genomes. Moreover, to be independent of the order in which genomes are added, we are interested in the average total size *f*_tot_(*m*) of the union for each possible species subset of size *m*, where 1 ≤ *m* ≤ *n*,

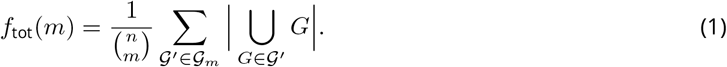

For convenience, we also define the average number *f*_new_(*m*) of new genes that are added when adding the *m*-th genome as:

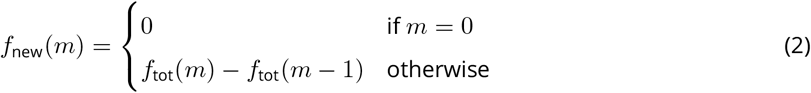

The function *f*_new_ is known to follow a power law distribution, referred to as Heaps’ law (Tettelin, Riley, et al., 2008), of the form *Km*^−*α*^ where *K* and *α* are positive, and the openness is defined based on the value of *α*. Since *f*_tot_ is defined as the cumulative sum over *f*_new_, we have that *f*_tot_ follows

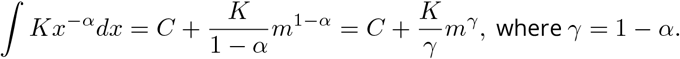

When *α <* 1, the function grows indefinitely as *m* goes to infinity, corresponding to an open pangenome. Conversely, when *α >* 1, *γ* becomes negative, such that *f*_tot_ rapidly approaches a constant value, *C*, resulting in a closed pangenome. The estimation of the pangenome’s openness is performed by fitting *Km*^−*α*^ on *f*_new_.

The original definition of openness provided by Tettelin, Riley, et al. (2008) differs from this, having *C* = 0 and thus 0 ≤ *γ* ≤ 1. This results in a contradiction, as it implies that *α* can never exceed one, suggesting that a closed pangenome is impossible.

To illustrate the concept of openness we show Figures 1 and 2 representing the fitting for some real genome data: *H. pylori* in red and *C. jejuni* in blue. Both datasets comprises 234 genomes each, have nearly identical genome lengths (*H. pylori*: 1.63Mbp; *C. jejuni*: 1.68Mbp), and contain a similar number of gene families (*H. pylori*: 1500; *C. jejuni*: 1700). Despite these similarities, *H. pylori* appears to be the more open of the two, with its pangenome containing two to three times more items than *C. jejuni*.

**Figure 1.**
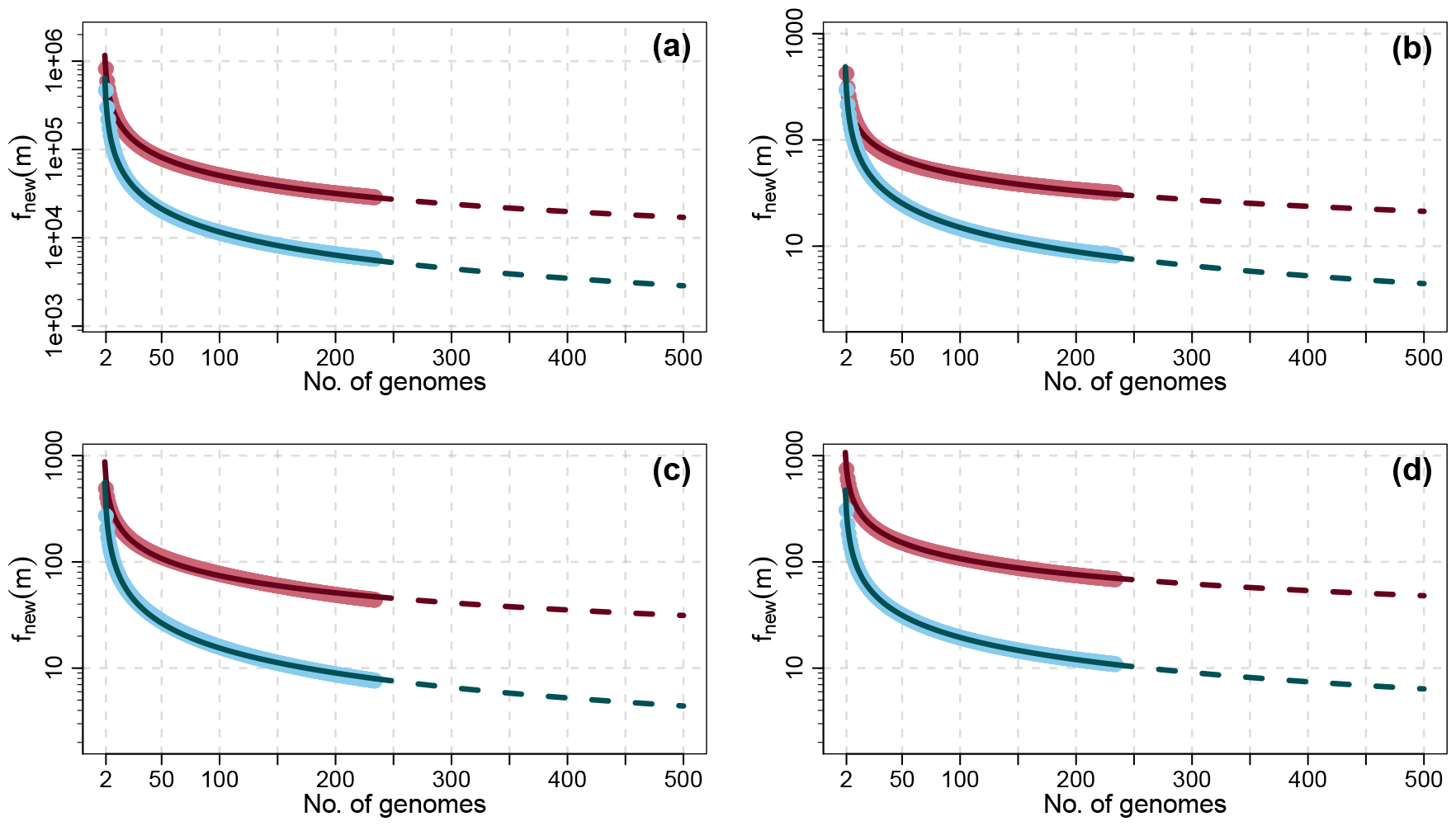
Comparison of the average number of new items *f*_new_(*m*) for 234 genomes of *Helicobacter pylori* (red) and for 234 genomes of *Campylobacter jejuni* (blue) using different tools: (a) Pangrowth, (b) Roary, (c) Pantools and (d) BPGA. The lines show the fitted functions *Km*^−*α*^.

**Figure 2.**
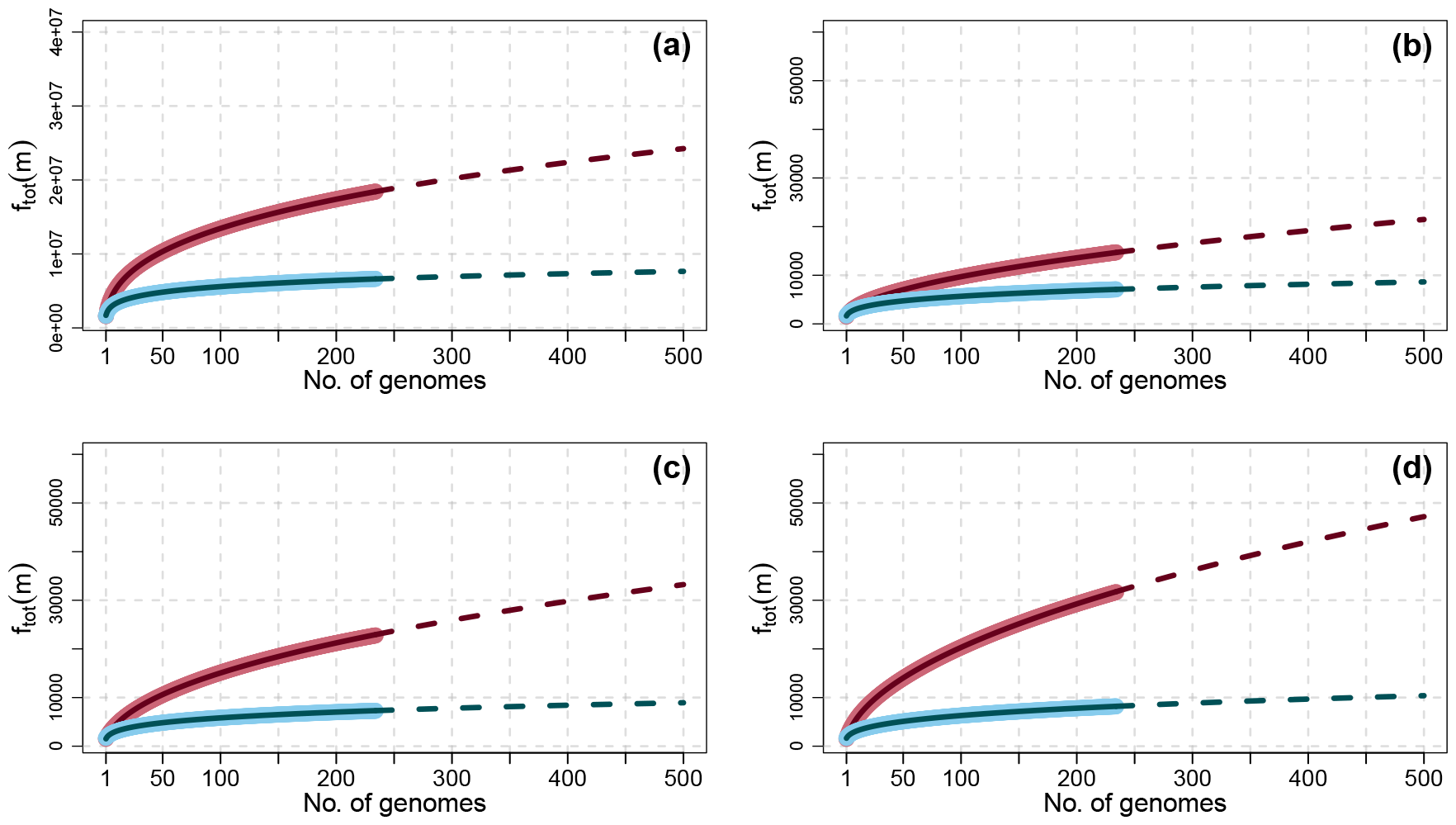
Comparison of the pangenome growth *f*_tot_(*m*) for 234 genomes of *Helicobacter pylori* (red) and for 234 genomes of *Campylobacter jejuni* (blue) using different tools: (a) Pangrowth, (b) Roary, (c) Pantools and (d) BPGA. The lines show the fitted functions 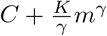.

In our analysis, *k*-mers were counted in their canonical form, i.e., a *k*-mer and its reverse complement are considered equivalent.

### Obtaining *f*_tot_ efficiently

The computation of *f*_tot_(*m*) requires taking the average of 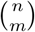 values when performed as formulated in Equation (1), making it prohibitive to compute even for relatively small *n*. The most common solution is to *approximate f*_tot_(*m*) selecting a subset of all the possible permutations of the order of the genomes, as suggested in (Tettelin, Riley, et al., 2008). Although this works well for gene-based approaches, it does not scale for *k*-mers. The reason is that the number of *k*-mers that can appear in a species can be two to four orders of magnitude higher than the number of genes (Table 1). This, in turn, slows down drastically the computation of the union of genomes which is linear in the number of items.

**Table 1.** Total number of distinct canonical *k*-mers found by Pangrowth and the total number of genes found by Roary (Page et al., 2015), Pantools (Sheikhizadeh et al., 2016) and BPGA (Chaudhari et al., 2016) for each species. Column “gene names” shows the total number of distinct gene names from the in silico annotation of Prokka (Seemann, 2014). In Pangrowth, the value chosen for *k* is 19 for all the species, except for *Buchnera aphidicola*, where *k* is 17.

| | species | $n$ | Pangrowth | Roary | Pantools | BPGA | gene names |
| --- | --- | --- | --- | --- | --- | --- | --- |
| 1 | *Bacillus cereus* | 83 | 47 161 713 | 36 794 | 37 532 | 43 014 | 2911 |
| 2 | *Buchnera aphidicola* | 35 | 12 341 389 | 13 441 | 13 240 | 13 571 | 295 |
| 3 | *Campylobacter jejuni* | 234 | 6 633 011 | 7097 | 7278 | 8196 | 902 |
| 4 | *Clostridium botulinum* | 50 | 32 016 575 | 27 528 | 25 948 | 28 982 | 2319 |
| 5 | *Coxiella burnetii* | 13 | 2 491 601 | 3433 | 2932 | 2767 | 242 |
| 6 | *Francisella tularensis* | 57 | 3 741 156 | 5137 | 3819 | 3726 | 767 |
| 7 | *Helicobacter pylori* | 234 | 18 430 113 | 14 708 | 22 774 | 31 672 | 790 |
| 8 | *Prochlorococcus marinus* | 9 | 12 502 284 | 13 088 | 12 612 | 13 300 | 421 |
| 9 | *Rhodopseudomonas palustris* | 8 | 21 985 762 | 15 135 | 14 678 | 15 841 | 788 |
| 10 | *Streptococcus pneumoniae* | 88 | 6 518 728 | 6375 | 6902 | 7059 | 818 |
| 11 | *Streptococcus pyogenes* | 247 | 7 420 059 | 7055 | 7138 | 7894 | 674 |
| 12 | *Yersinia pestis* | 56 | 4 917 598 | 6087 | 5612 | 5362 | 889 |

Here we describe a more efficient, *exact* way to calculate *f*_tot_(*m*), without computing 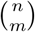 subsets for each average. This method has been known for several decades in the field of ecology (Heck et al., 1975) but it seems to have gone unnoticed in the fields of pangenomics and linguistics (Chacoma and Zanette, 2020). Let *h*(*i*) be the number of items that occur in exactly *i* of the *n* input genomes, 1 ≤ *i* ≤ *n*. We consider the contribution of each item of multiplicity *i* based on the number of subsets *𝒢*^*′*^ ∈ *𝒢*_*m*_ in which the item will be counted. This can be done by subtracting from the number of subsets in 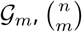, the number of subsets that *do not* contain an item of multiplicity *i*, 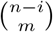. We can then rewrite Equation (1) as follows:

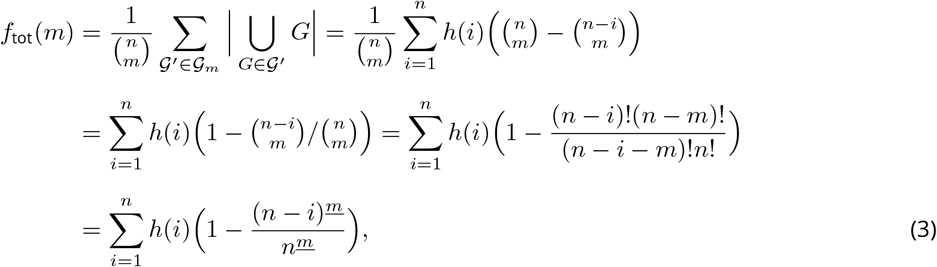

where *n*^*<u>m</u>*^ is the falling factorial defined as *n*^*<u>m</u>*^ = *n*(*n* − 1)(*n* − 2) *…* (*n* − *m* + 1). Observe that *n*^*<u>m</u>*^ = 0 if *m > n*, since there is a zero in the product.

### Estimating the pangenome openness

Our procedure for the estimation of the pangenome openness is divided into three steps. First, the values *h*(*i*) must be computed or provided. As a matter of computing, they can be extracted directly from the pan-matrix by counting the number of rows that sum up to *i*. More efficient solutions can be considered based on the item of choice as shown in the following subsection. In the second step, *f*_tot_ is calculated according to Equation (3). We can rewrite Equation (3) as 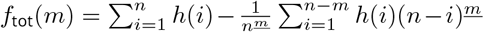 and compute (*n* −*i*)^*<u>j</u>*<u>+1</u>^ = (*n* − *i* − *j*)(*n* − *i*)^*<u>j</u>*^, where (*n* − *i*)^<u>1</u>^ = (*n* − *i*). In practice, we can store the values (*n* − *i*)^*<u>j</u>*^ reducing the time complexity of the calculation of *f*_tot_ from Θ(*n*^3^) to Θ(*n*^2^), with the same space complexity, Θ(*n*). Lastly, *f*_new_ is computed from *f*_tot_ using Equation (2), and the function *Km*^−*α*^ is fitted on *f*_new_, estimating the parameter *α* for the pangenome openness.

### Implementation using *k*-mers as items

We have implemented the estimation of the pangenome openness that uses *k*-mers as items in a software called Pangrowth. The code is available as a Gitlab project^1^.

For the computation of *h*(*i*) we modified the *k*-mer counting tool yak (Cheng et al., 2021). Since storing *k*-mers in the pan-matrix is not efficient in terms of memory, yak uses multiple hash tables for different suffixes of *k*-mers, reducing access time and enabling parallelization. To prevent counting the same *k*-mer twice in the same genome, we associate a variable to each *k*-mer representing the last genome it was found in. Only if this is different from the genome that is currently read, the counter is incremented. The histograms for *H. pylori* and *C. jejuni* are shown in Figures 3(a) and 3(b), respectively. Next, we compute *f*_tot_ following the approach outlined in the ‘Obtaining *f*_tot_ efficiently’ subsection, determine *f*_new_ and estimate *α*. Figure 1 reports *f*_new_ computed on the two datasets with their fitting for each of the four tools under study.

**Figure 3.**
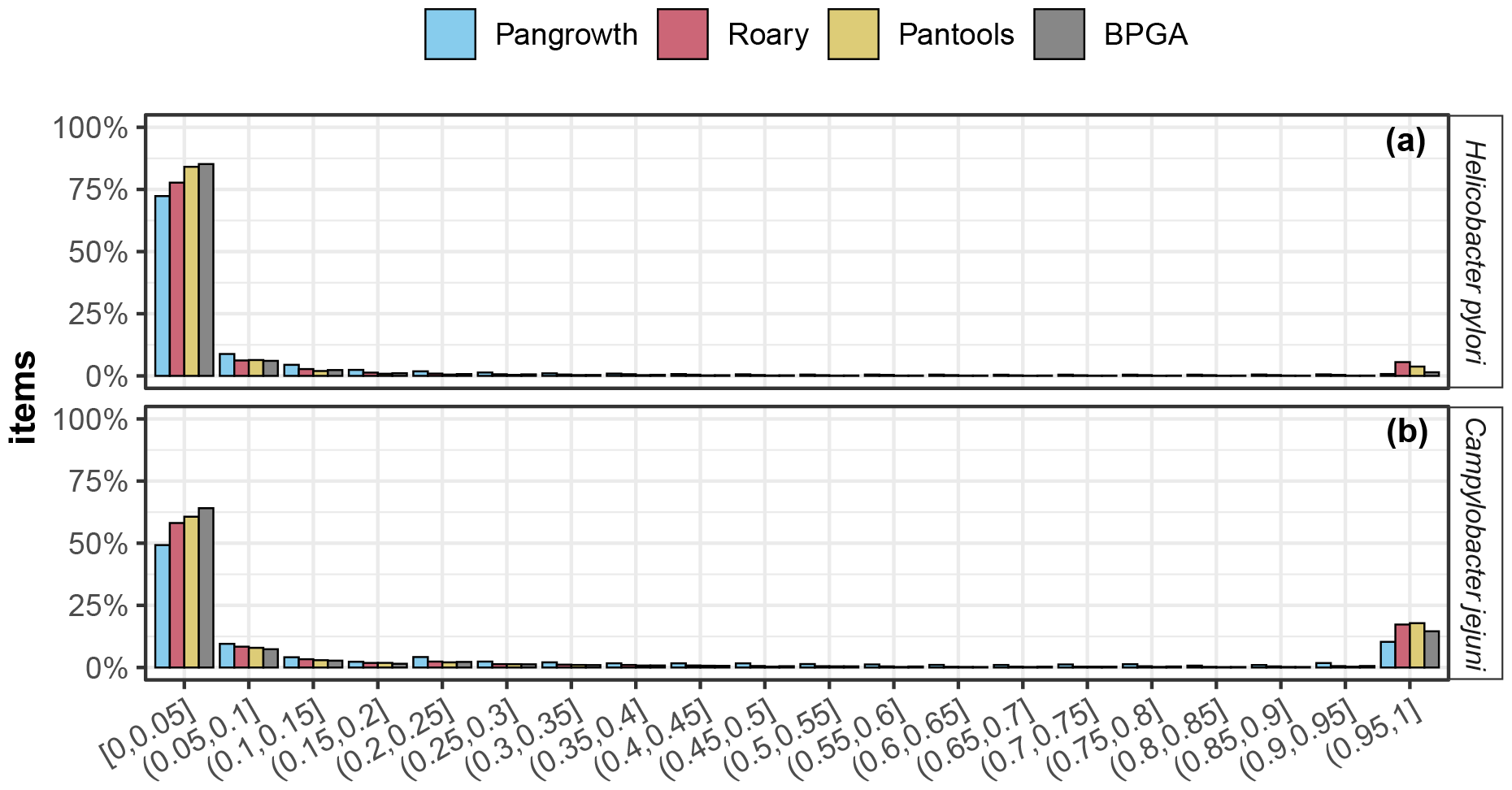
Percentage of items in histogram *h*(*i*) in (a) 234 genomes of *Helicobacter pylori* and in (b) 234 genomes of *Campylobacter jejuni* for the tools Pangrowth, Roary, Pantools and BPGA. The *y*-axis displays the normalized values of the histogram, while the *x*-axis represents the percentage of genomes in which the items appear, with values binned together in 5% increment, for visual clarity.

## Results

We analysed twelve bacterial pangenomes with the three gene-based approaches, Roary (Page et al., 2015), Pantools (Sheikhizadeh et al., 2016) and BPGA (Chaudhari et al., 2016), and with our *k*-mer-based approach, Pangrowth. To rule out sampling as a possible source of difference in the results, we used Equation (3) on the pan-matrix generated by each gene-based tool. Pantools is capable of directly producing the pan-matrix, thus avoiding the sampling across multiple genome orders. However, Roary and BPGA do not offer this functionality. We altered Roary’s code to end the process right after the pan-matrix is generated. Conversely, BPGA requires a minimum of 20 permutations of genome orders and only after completing this task it outputs the pan-matrix. Since BPGA comes in binary format, we could not modified it.

Genomes of each species were downloaded from NCBI RefSeq with the filter “Assembly level: Complete genome”. Annotations have been carried out by Prokka (Seemann, 2014) version 1.14.6 with standard parameters. The pipeline was wrapped in a Snakemake workflow for reproducibility and parallelization with the exception of BPGA, for which we provide a separate bash script. The number of genomes for each species can be found in Table 1 while the value of *α* for each tool is reported in Table 2. Both the Supplementary Material and the data with scripts can be accessed at Zenodo^2^.

**Table 2.** Values of *K* and *α* for Pangrowth, Roary, Pantools and BPGA for each species. The values of *K* were approximated to the nearest integer.

| species | $K$ | | | | $\alpha$ | | | |
| --- | --- | --- | --- | --- | --- | --- | --- | --- |
|  | Pangrowth | Roary | Pantools | BPGA | Pangrowth | Roary | Pantools | BPGA |
| 1 *Bacillus cereus* | 5 756 858 | 3354 | 3503 | 4215 | 0.77 | 0.68 | 0.68 | 0.69 |
| 2 *Buchnera aphidicola* | 755 172 | 642 | 651 | 653 | 0.30 | 0.20 | 0.21 | 0.20 |
| 3 *Campylobacter jejuni* | 640 315 | 487 | 560 | 473 | 0.87 | 0.76 | 0.78 | 0.69 |
| 4 *Clostridium botulinum* | 4 120 907 | 3109 | 3218 | 3496 | 0.71 | 0.66 | 0.70 | 0.68 |
| 5 *Coxiella burnetii* | 200 334 | 499 | 374 | 288 | 0.91 | 0.87 | 0.92 | 0.83 |
| 6 *Francisella tularensis* | 301 531 | 412 | 272 | 211 | 0.76 | 0.71 | 0.73 | 0.64 |
| 7 *Helicobacter pylori* | 1 160 492 | 442 | 873 | 1069 | 0.68 | 0.49 | 0.54 | 0.50 |
| 8 *Prochlorococcus marinus* | 1 958 786 | 2206 | 2212 | 2228 | 0.24 | 0.29 | 0.32 | 0.29 |
| 9 *Rhodopseudomonas palustris* | 5 139 651 | 2992 | 2874 | 3312 | 0.53 | 0.49 | 0.49 | 0.51 |
| 10 *Streptococcus pneumoniae* | 549 066 | 516 | 651 | 541 | 0.74 | 0.73 | 0.77 | 0.69 |
| 11 *Streptococcus pyogenes* | 472 284 | 570 | 457 | 418 | 0.74 | 0.81 | 0.74 | 0.68 |
| 12 *Yersinia pestis* | 70 781 | 381 | 172 | 156 | 0.85 | 0.87 | 0.62 | 0.64 |

### Choice of parameters

Roary was run using its default parameters. For Pantools, the mandatory relaxation parameter, which ranges from one to eight, was set to one. This parameter controls how similar two gene sequences must be to be considered part of the same gene family, with a value of one requiring the sequences to be closely related^3^. In BPGA, we used the default clustering method, USEARCH, since CD-HIT and MCL were already employed in Roary and Pantools, thereby adding a new clustering method to the list. The sequence similarity threshold was set to 0.95. We note that a subsequent run with a threshold of 0.97 did not lead to significant changes in the reported values of *α*.

In our method, the only parameter is *k* for which we used a similar heuristic to what was proposed in (Sheikhizadeh Anari et al., 2018): Given the length of the genome, the value of *k* is chosen to be the smallest that ensures the probability of a random *k*-mer to occur in a genome is below 0.0001, rounded to the next odd integer. On the considered datasets, this heuristic suggests *k* = 19 for almost all species, since the scale of the size of bacterial genomes is similar, with the exception of *Buchnera aphidicola*, where *k* = 17 is proposed due to its smaller genome size. Even though inversions and other rearrangements that affect the direction of substrings within the genome can be accounted as variability between genomes, we are more interested in new items (genes or *k*-mers). For this reason we chose to look at canonical *k*-mers and only *k*-mers that span rearrangement breakpoints are affected.

### Fitting Heaps’ law

The reported values for *α* were determined by applying a linear model to the log-log scale of *f*_new_. Figures S1, S2, S3 and S4 illustrate the adjusted *R*^2^ values for the fitting of Heaps’ law across various species at different starting positions 2 ≤ *m*_0_ ≤ 10. This is shown for our *k*-mer-based approach as well as for the three gene-based approaches. In the *k*-mer-based approach, *Yersinia pestis* is the most influenced by the starting position of the fit, initially not being very well described by Heaps’ law (*R*^2^ = 0.91) and increasingly fitting better with higher starting positions (*R*^2^ *>* 0.99 for *m*_0_ ≥ 6). Similarly, *Rhodopseudomonas palustris* shows an *R*^2^ for *m*_0_ = 2 between 0.92 and 0.95 for the gene-based approaches, that also increases with higher *m*_0_. For the remaining species, and across all tools, the power law seems to consistently provide a good fit (*R*^2^ = 0.96 for *m*_0_ = 2 and *R*^2^ *>* 0.99 for *m*_0_ ≥ 6), with the value of *α* not significantly altered when *m*_0_ is changed (Figures S5, S6, S7 and S8).

That the fit improves when *m*_0_ is increased is in line with the pattern commonly observed in other phenomena where only the tail of the function follows the power law (Clauset et al., 2009). Depending on the interest, the fit can be adjusted for the tail, if estimating the average number of newly discovered items for newly sequenced genomes is the goal, or the fit can be done as early as possible, to estimate the species’ openness. For example, in Tettelin, Riley, et al. (2008) they start at *m*_0_ = 3, while most tools begin at *m*_0_ = 2 (Sheikhizadeh et al., 2016; Snipen and Liland, 2015). In this paper, we are interested in the latter approach. Therefore, unless otherwise stated, all the reported values of *α* are fitted with *m*_0_ = 2.

### Overall comparison

Figure 4 displays a pairplot comparing the *α* values across the tools considered. The gene-based tools show consistent results, especially between Pantools and BPGA, where they report nearly identical values. Similarly, Roary and BPGA, as well as Roary and Pantools, agree on their values, except for *Y. pestis*. In this case, Pantools and BPGA converge on one value around 0.630, while Roary reports 0.868. Our *k*-mer-based approach, Pangrowth, shows strong correlation with the gene-based tools, with a Pearson correlation coefficient *ρ >* 0.92, and reports similar values of *α* with respect to all three gene-based tools.

**Figure 4.**
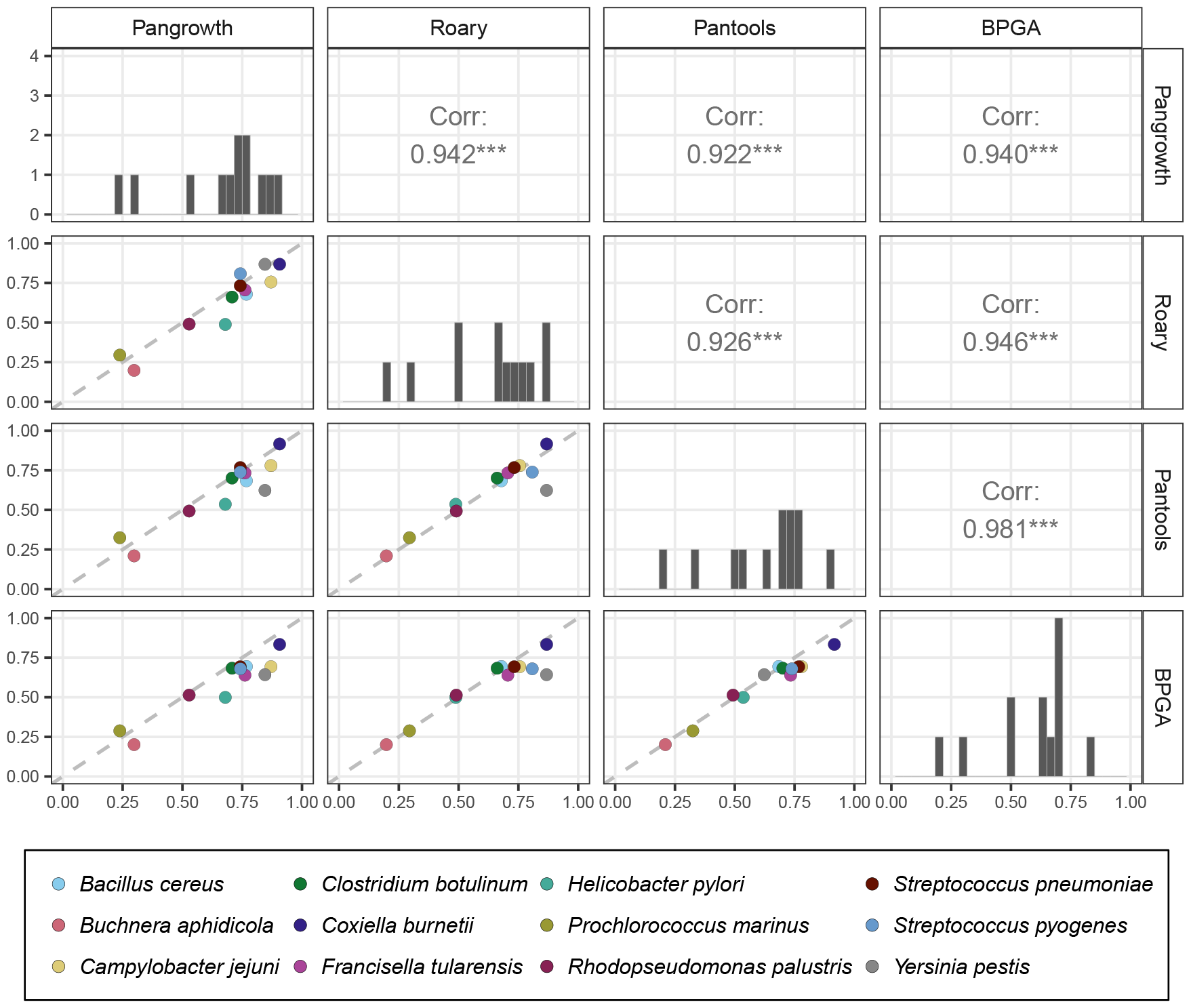
Pairplot illustrating the comparison of estimated openness values across the 12 bacterial species and the four tools. The lower triangle shows the comparison of *α* values for each species, with the diagonal dashed line representing the equality. The upper triangle depicts the Pearson correlation coefficient, where ‘***’ signifies a p-value *<* 0.001. The diagonal line contains the distribution of the *α* values with the *y*-axis ranging from 0 to 4.

To estimate how sensible the openness is to the choice of *k*, we computed *α* for different values of *k* (Figure 5). While more open species appear to be more influenced by *k*, the relative order of the species stabilizes for *k* ≥ 21.

**Figure 5.**
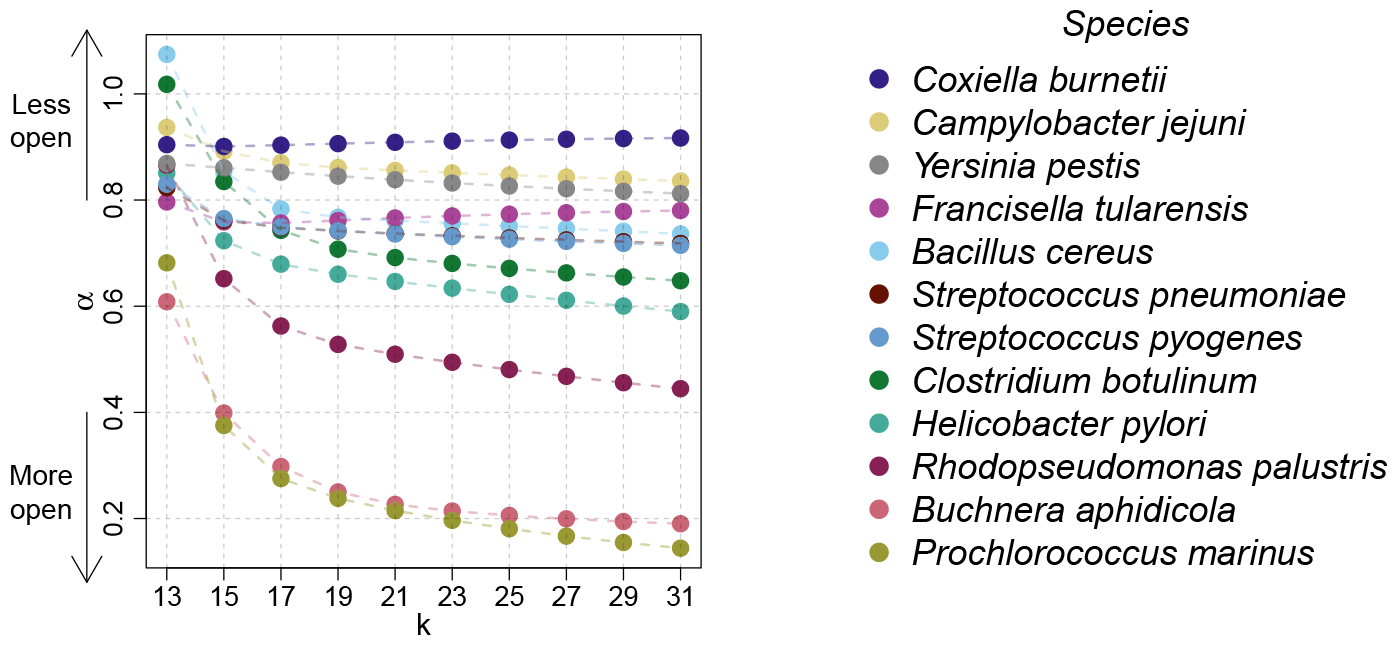
Values of *α* for different choices of *k*.

We compared the running time and space consumption of our tool against the three gene-based tools and show the results in Figure 6. All computations and measurements were conducted on our server machine, equipped with 28 Intel Xeon Processors (2.6 GHz) and 64 GB of RAM. Every tool had 28 threads at disposal. In terms of running times, our method is one to three orders of magnitude faster then any of the gene-based methods, with Prokka annotations given. Note that both Roary and BPGA have potential for speed improvements. They could be optimized by focusing solely on generating the pan-matrix or, more effectively, a histogram similar to that of Pangrowth. In terms of memory, Pangrowth uses less than 1 GB of memory for each of the species, similarly to Roary and BPGA, while Pantools requires more memory, probably due to the vast features of the tool. Prokka consistently uses less than 100 MB of memory for each species and takes between 4 minutes and 1 hour to annotate, depending on the dataset, running on a single thread.

**Figure 6.**
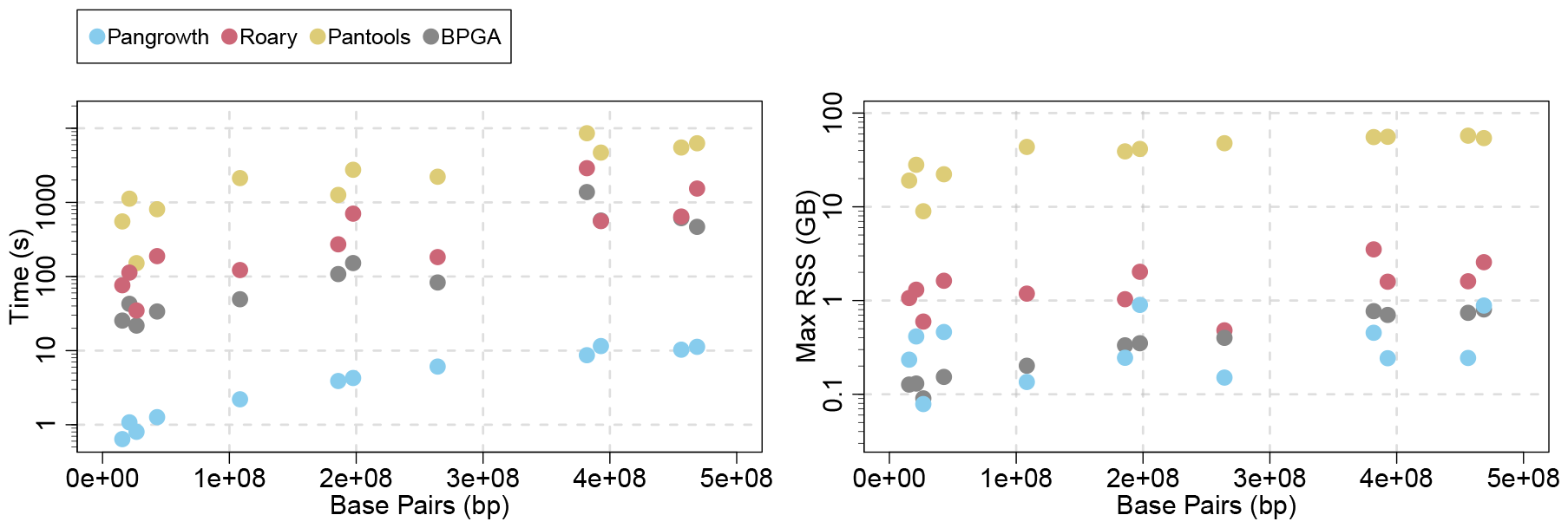
Wall clock time (left) and Maximum Resident Set Size (right) measured for each species and across all tools.

To evaluate Pangrowth’s scalability, we conducted a series of benchmarks on a set of 8000 *Escherichia coli* genomes, incrementally increasing the sample size across tests. Due to the low resources needed, these tests were run on a portable computer furnished with 12 Intel Core i7-8750H processors (2.20 GHz) and 16 GB of RAM. The wall clock times for these tests are presented in Figure 7 for *k* = 19. Although the construction of the pangenome growth has a complexity quadratic in the number of genomes, the pangenome growth of a histogram of 8000 *E. coli* genomes can be computed in under a second, requiring below 4 MB of memory. This indicates that the bottleneck is in the calculation of the histogram, rather than in the growth computation. Even so, this step runs linear in practice with respect to the number of genomes. On average, the calculation took approximately 19 minutes for 8000 genomes and consumed around 3.5 GB of memory.

**Figure 7.**
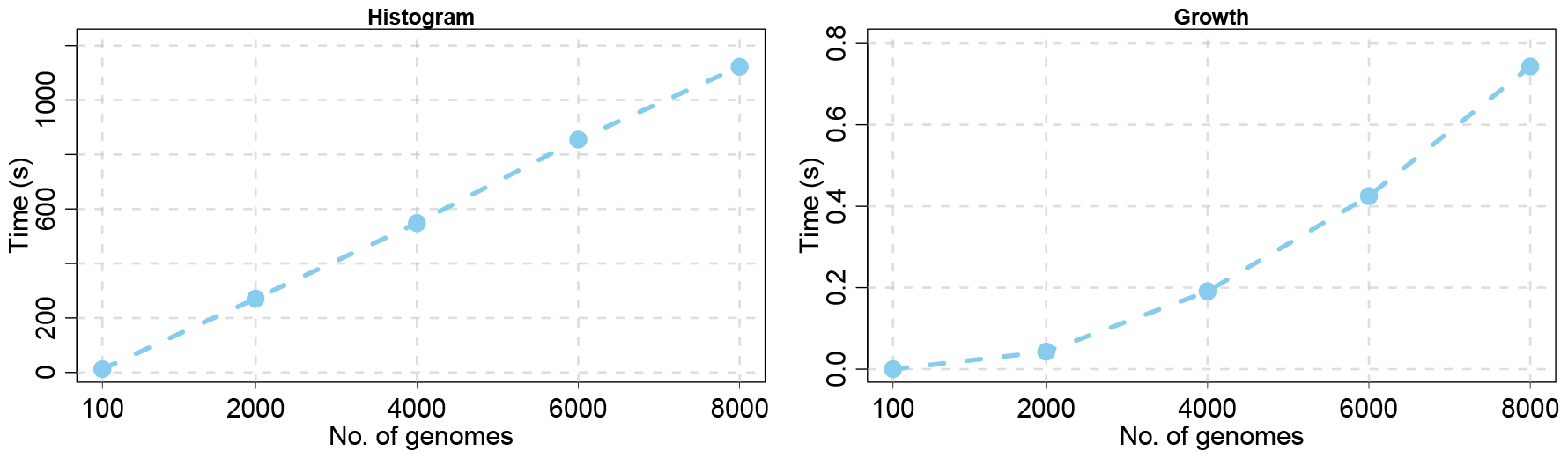
User time measured by the /usr/bin/time command, across different sample sizes of 8000 genomes of *Escherichia coli*. The measurements are reported separately for the two distinct stages of Pangrowth: (left) the generation of the histogram *h*, and (right) the subsequent calculation of the pangenome growth based on this histogram. Each point represents the average over 10 runs.

Overall, results show similar trends in the estimated openness over multiple species between the *k*-mer-based and the gene-based methods. Combined with the absence of outliers and the fast computation time, they support the use of *k*-mers as a reasonable and practical alternative to genes in the estimation of the openness.

### Closeness

In our analysis, we did not deliberately exclude closed pangenomes but found none that met the criteria for being closed as defined by Tettelin, Riley, et al. (2008). While other studies have reported closed pangenomes, it is important to recognize the complexity in comparing these findings, as numerous variables can influence whether a pangenome is classified as closed or not.

Firstly, the use of various fitting methods, such as different starting points, fitting over all points or over the median, combined with a lack of goodness-of-fit, can lead to different conclusions.

Secondly, certain species were initially considered closed in the early stages of pangenomics due to limited genome availability. For example, Tettelin, Riley, et al. (2008) identified *Staphylococcus aureus* as having a closed pangenome. However, this was later revised to open by Bosi et al. (2016).

Lastly, some pangenomes are reported as closed with *α* values just below one. For instance, the study by Argemi, Matelska, et al. (2018) declared the pangenome of *Staphylococcus lugdunensis* as closed with *α* = 0.96. Even though their method of fitting a power law over *f*_tot_ and reporting *α* as 1 − *γ* cannot return a closed pangenome, we obtain the same *α* by taking the pangenome growth they provided, compute *f*_new_, and fit *α* = 0.91 (*R*^2^ = 0.96). Using Pangrowth on this set of genomes we obtain *α* = 0.92 (*R*^2^ = 0.97). Despite observing an *α* considerably smaller than 1, *S. lugdunensis* was classified by Argemi, Hansmann, et al. (2019) as closed also due the high similarity of their genomes. Following the same rationale, *Yersinia pestis* can be classified as closed, which is in line with other studies (Cui and Song, 2016).

We found that Coccolitovirus (*Phycodnaviridae*) shows a mathematically closed pangenome. Using the panmatrix reported by Lobb et al. (2023), we computed the pangenome growth and calculated *α*. As reported in their manuscript, Coccolitovirus is closed with *α* = 1.21 (*R*^2^ = 1.00) (in their manuscript they report 1.22 based on 1000 permutations). Using Pangrowth on the same genomes of Coccolitovirus we obtain even *α* = 1.94 (*R*^2^ = 0.94).

## Conclusion

The use of *k*-mers to process pangenomes is common practice. In fact, *k*-mers are almost ubiquitous when processing and analysing genetic sequences. The main focus of this manuscript is the comparison of genes and *k*-mers when computing the pangenome openness. We showed that the values of *α* resulting from the *k*-mer-based approach in bacterial species is similar to the results obtained with a gene-based approach.

One of our motivations for the use of *k*-mers is the possibility to assess the openness also for non-bacterial species. The concept of openness can be naturally extended to non-bacterial genomes as a measure of the richness of items present in a species. While bacterial genomes are notoriously composed mostly of protein coding regions, this is not true for eukaryotic genomes. Given the significant role that non-coding regions play in eukaryotic genomes, sequence-based approaches such as *k*-mer analysis may be more effective in capturing the full genomic content.

To test our method with a non-bacterial dataset, we applied our tool to 100 randomly selected human genomes from the 1000 Genomes Project (1000 Genomes Project Consortium, 2015), containing only autosomes. The construction of the histogram took around 1 hour and 15 minutes on our server, while the time for the construction of the pangenome growth from the histogram was around 10 milliseconds. For this dataset we observed *α* = 0.789.

Our experiments show that different gene homology definitions can vary the prediction of the openness. Our *k*-mer approach does not require gene homology but it still needs a proper choice of *k*. However, except for small values of *k* where almost all *k*-mers appear, the values of *α* are not very sensible to the choice of *k*.

Another common pangenome analysis is the identification of the pangenomic core. Equation (3) can be promptly modified to estimate the size of the core. Similarly to the computation of *f*_tot_, the core estimation requires taking the average number of shared items. The only required modification in *f*_tot_ is the use of the intersection of each genome instead of the union. We can also relax the definition of core items to count the items present in a subset of the set of genomes, e.g., items present in 90% of the genomes.

## Acknowledgements

We would like to thank the reviewers for their careful reading and helpful suggestions to improve the manuscript.

## Fundings

This project received funding from the European Union’s Horizon 2020 research and innovation programme under the Marie Skłodowska-Curie grant agreement No 956229. It was also supported by the BMBF-funded de.NBI Cloud within the German Network for Bioinformatics Infrastructure (de.NBI) (031A532B, 031A533A, 031A533B, 031A534A, 031A535A, 031A537A, 031A537B, 031A537C, 031A537D, 031A538A).

## Conflict of interest disclosure

The authors declare that they comply with the PCI rule of having no financial conflicts of interest in relation to the content of the article.

## Footnotes

1 https://gitlab.ub.uni-bielefeld.de/gi/pangrowth

2 Supplementary Material: https://zenodo.org/record/8233908. Data and scripts: https://zenodo.org/record/8256094

3 We communicated with the authors of Pantools after an initial discrepancy in the results. The divergence arose for two reasons: we set the gene homology parameter too high (relaxation was set to 4 instead of 1), and because Pantools fits Heaps’ law across all data points obtained from the permutations of the order of genomes, rather than fitting it over the average or the median.

## Notes

https://gitlab.ub.uni-bielefeld.de/gi/pangrowth

https://zenodo.org/record/8233908

https://zenodo.org/record/8256094

### Competing Interest Statement

The authors have declared no competing interest.

### Summary of Updates

Two new datasets were added to the comparison and a new subsection called Openness was added to describe them. Some general changes in the structure of the paper. Rerun of timing for gene-based tools based on the pan-matrix. New plot for timing of pangrowth of E. coli.

