## Supplementary File for "Revisiting pangenome openness with *k*-mers"

### Supplementary Material

#### Contents

**S1 Fitting Heaps' law** 3

**S2 Pangrowth** 7

**S3 Roary** 11

**S4 Pantools** 15

**S5 BPGA** 19

**S6 Histograms** 23

### S1 Fitting Heaps' law

**Figure S1.** Adjusted $R^2$ values for the fitting of **Pangrowth**, over different starting points. A minimum of 6 genomes was required to fit the data.

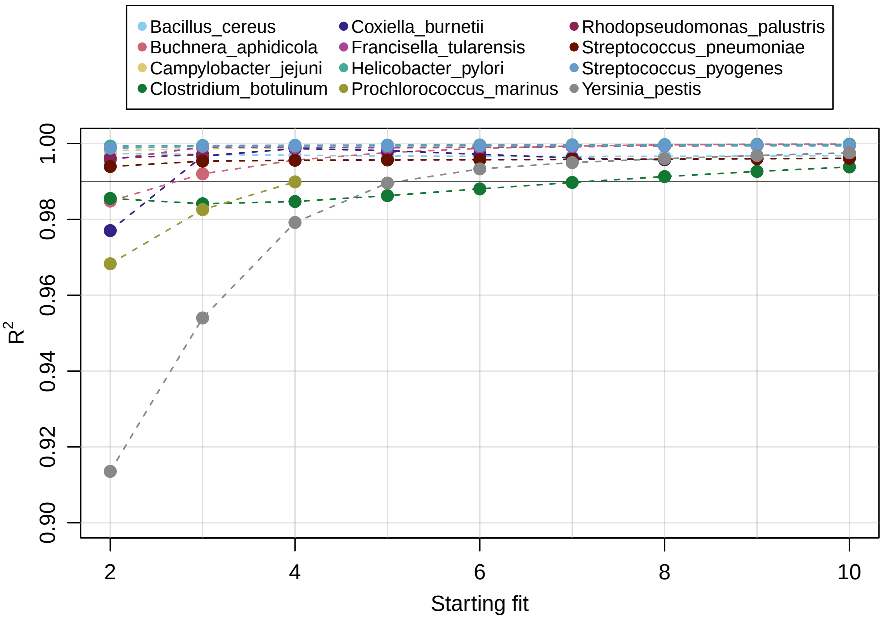


**Figure S2.** Adjusted $R^2$ values for the fitting of **Roary**, over different starting points. A minimum of 6 genomes was required to fit the data.

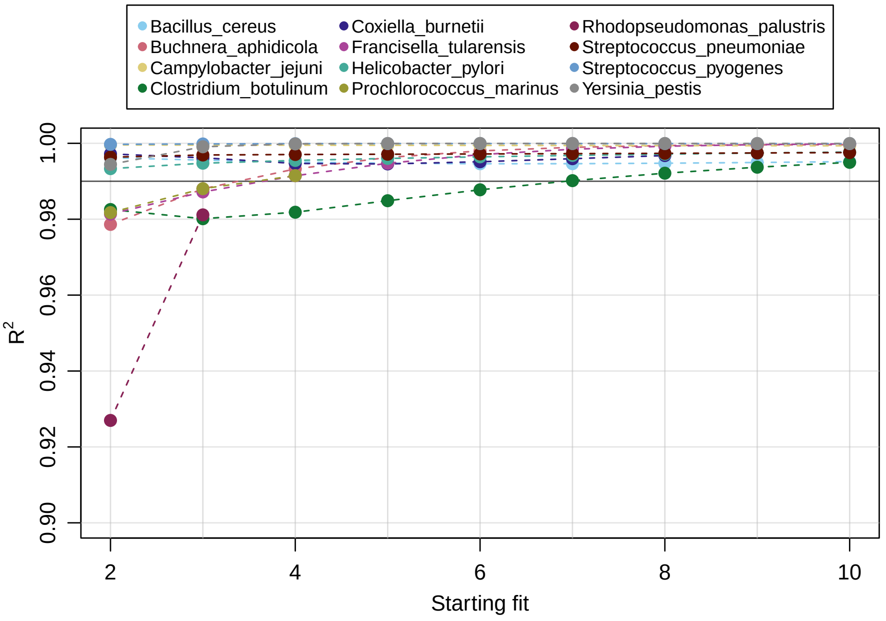

**Figure S3.** Adjusted $R^2$ values for the fitting of **Pantools**, over different starting points. A minimum of 6 genomes was required to fit the data.

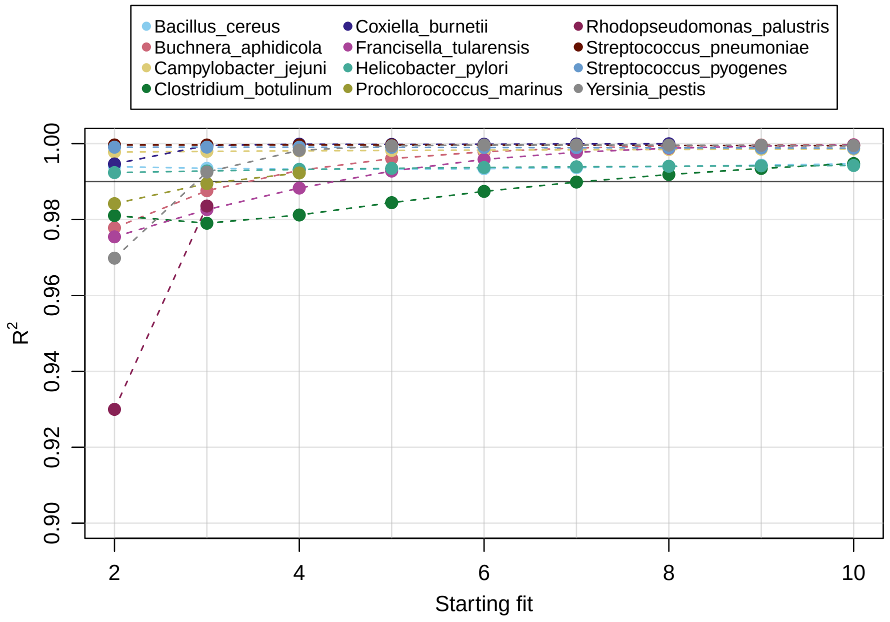


**Figure S4.** Adjusted $R^2$ values for the fitting of **BPGA**, over different starting points. A minimum of 6 genomes was required to fit the data.

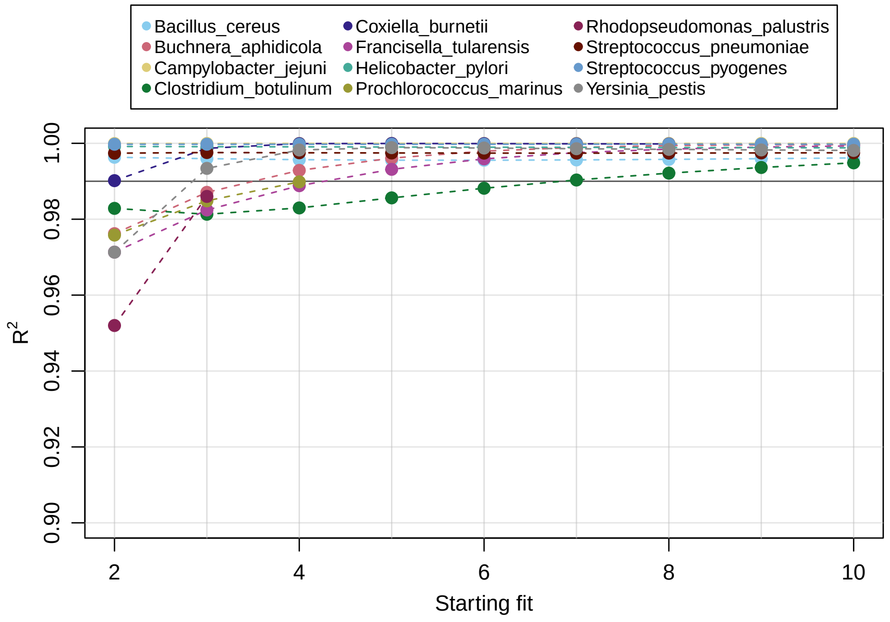

**Figure S5.** Estimated value of $\alpha$ for **Pangrowth**, over different starting points. A minimum of 6 genomes was required to fit the data.

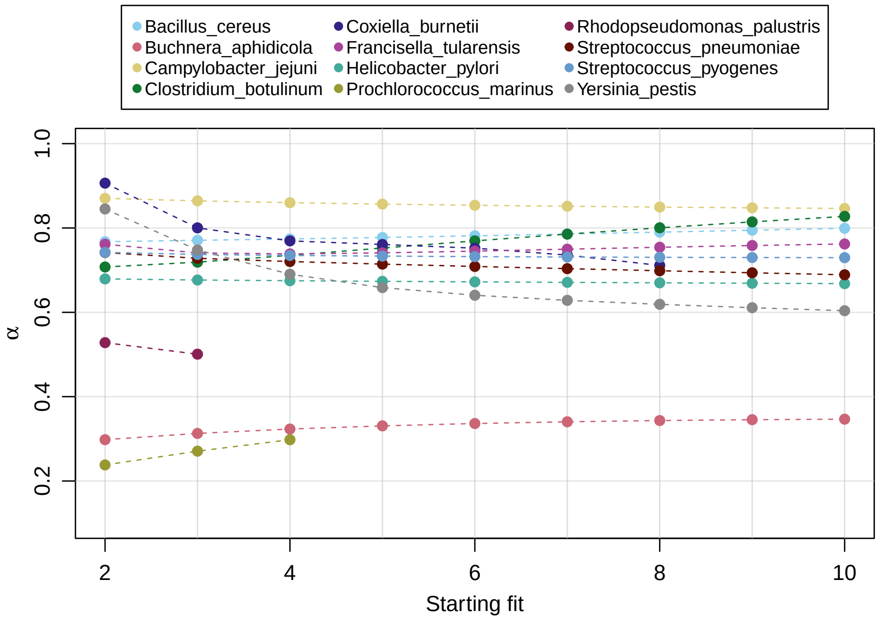


**Figure S6.** Estimated value of $\alpha$ for **Roary**, over different starting points. A minimum of 6 genomes was required to fit the data.

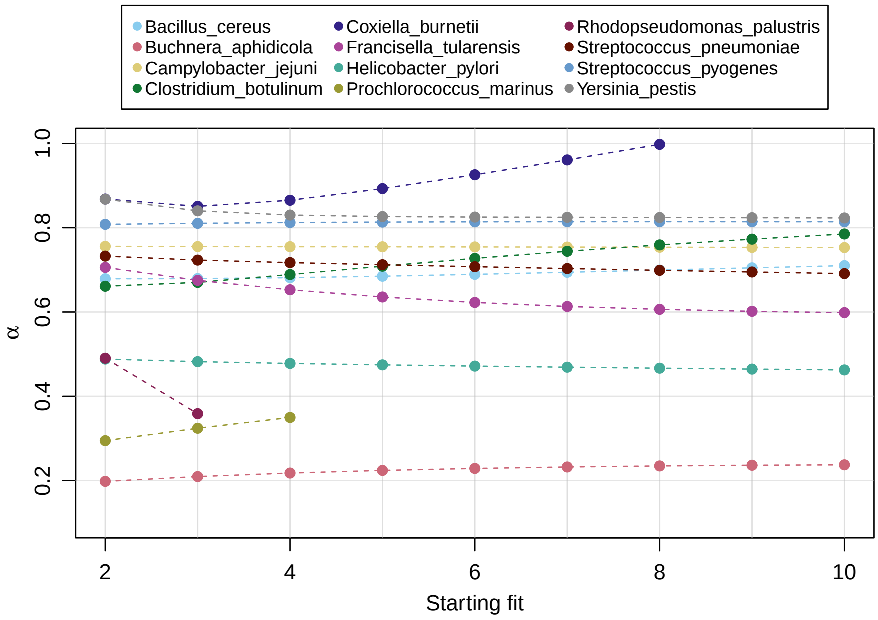

**Figure S7.** Estimated value of $\alpha$ for **Pantools**, over different starting points. A minimum of 6 genomes was required to fit the data.

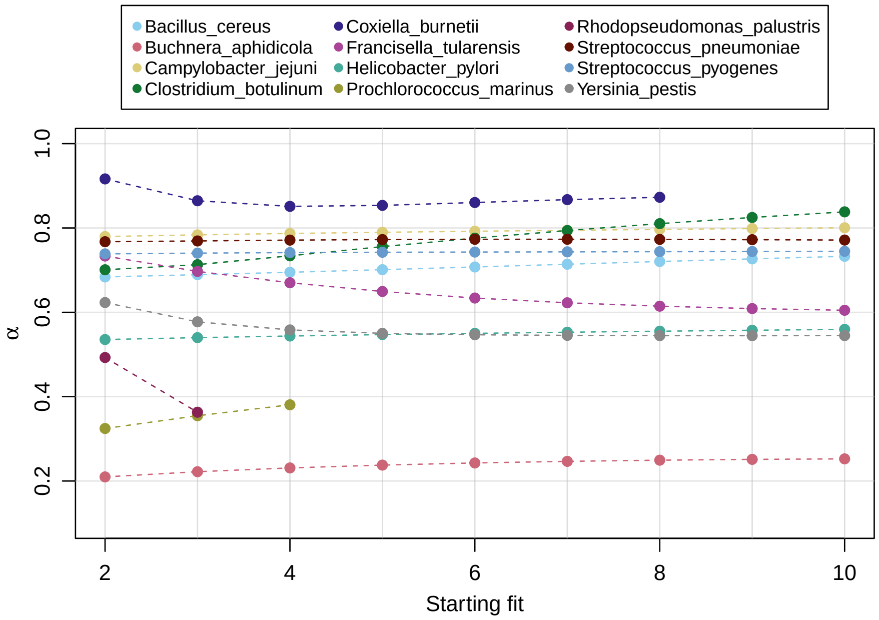


**Figure S8.** Estimated value of $\alpha$ for **BPGA**, over different starting points. A minimum of 6 genomes was required to fit the data.

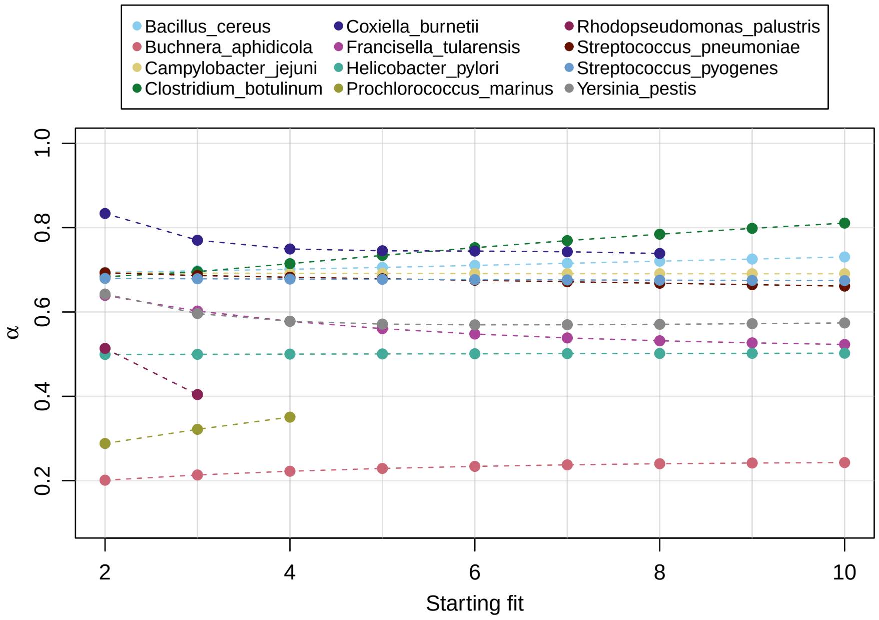

#### S2 Pangrowth

**Figure S9.** Average growth of *Bacillus cereus*.

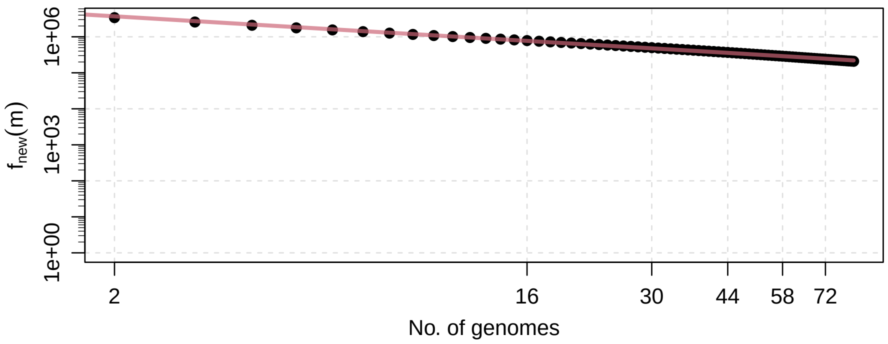


**Figure S10.** Average growth of *Buchnera aphidicola*.

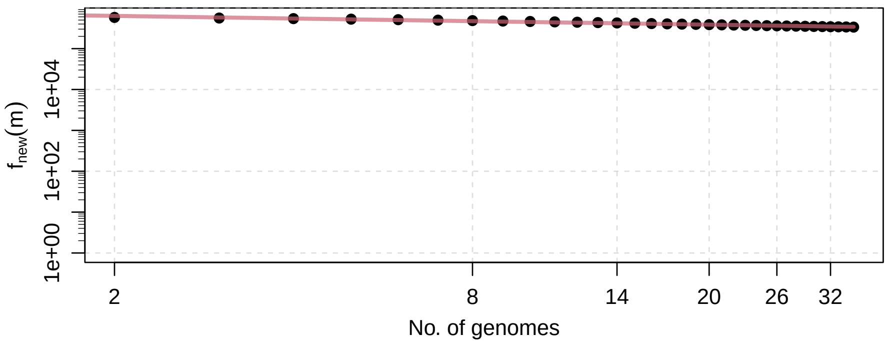


**Figure S11.** Average growth of *Campylobacter jejuni*.

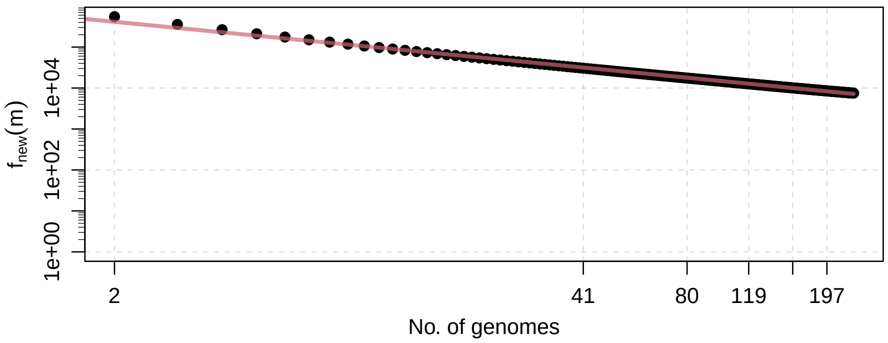

**Figure S12.** Average growth of *Clostridium botulinum*.

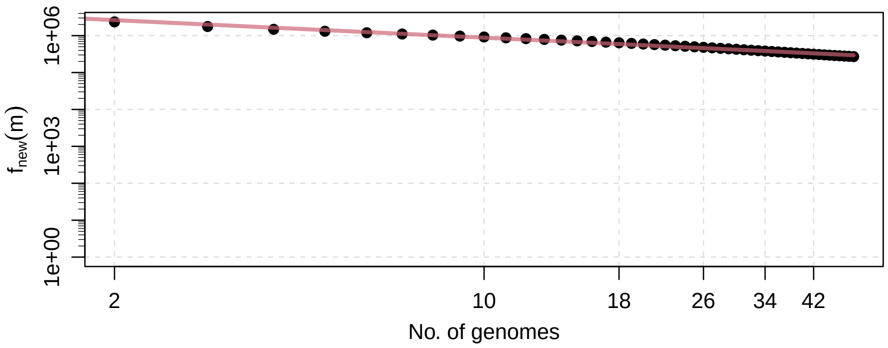


**Figure S13.** Average growth of *Coxiella burnetii*.

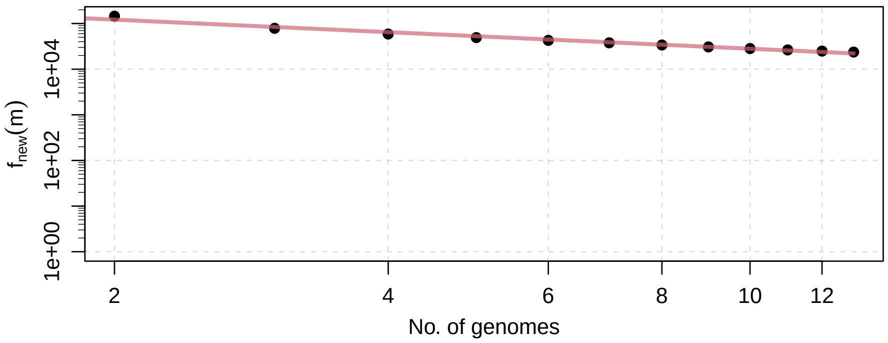


**Figure S14.** Average growth of *Francisella tularensis*.

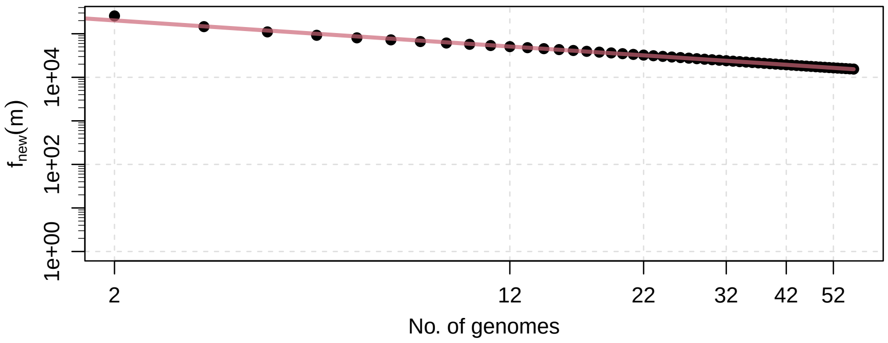

**Figure S15.** Average growth of *Helicobacter pylori*.

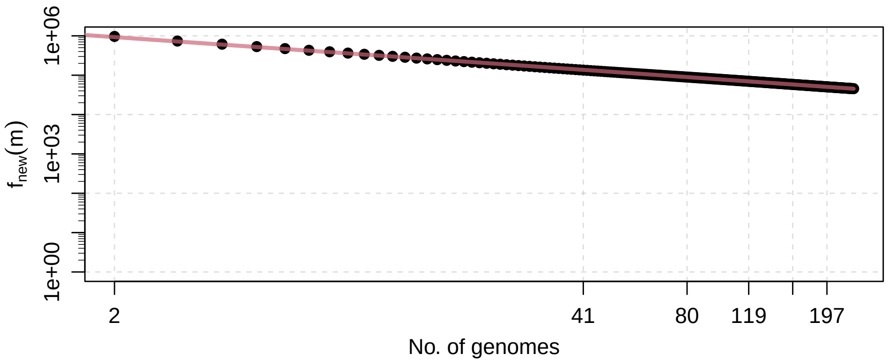


**Figure S16.** Average growth of *Prochlorococcus marinus*.

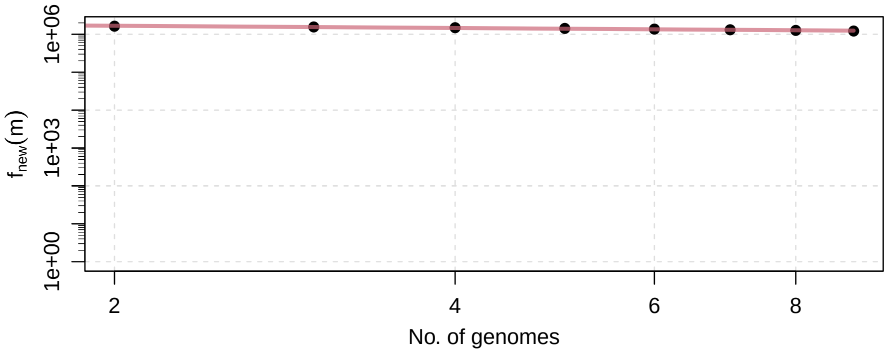


**Figure S17.** Average growth of *Rhodopseudomonas palustris*.

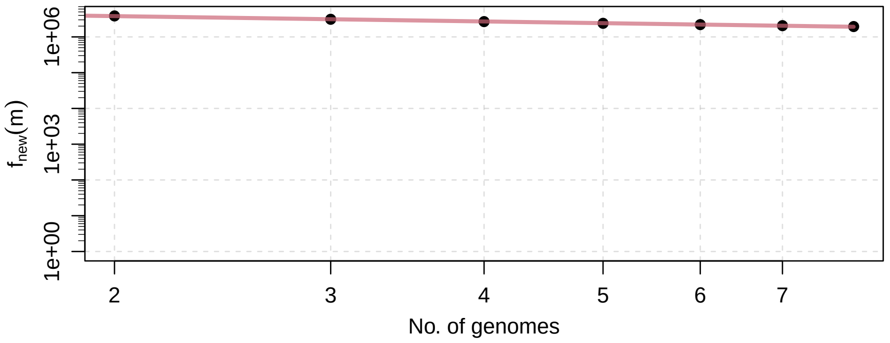

**Figure S18.** Average growth of *Streptococcus pneumoniae*.

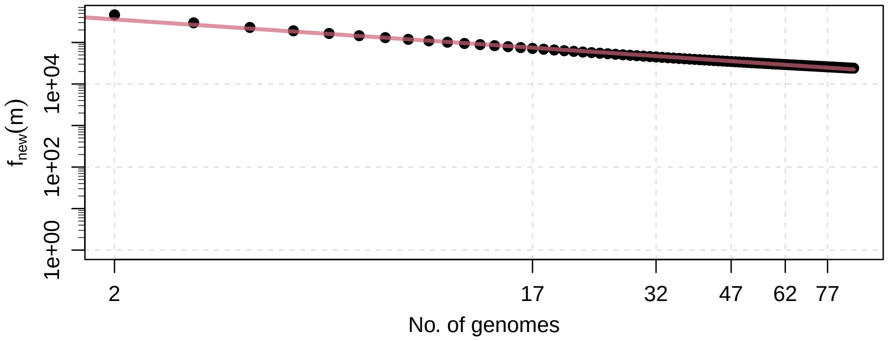


**Figure S19.** Average growth of *Streptococcus pyogenes*.

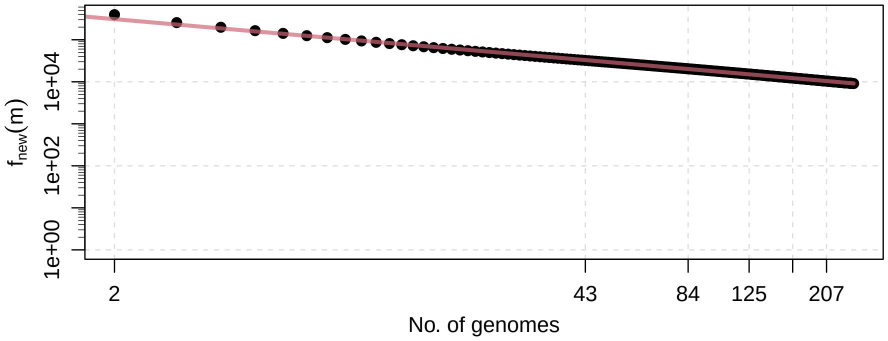


**Figure S20.** Average growth of *Yersinia pestis*.

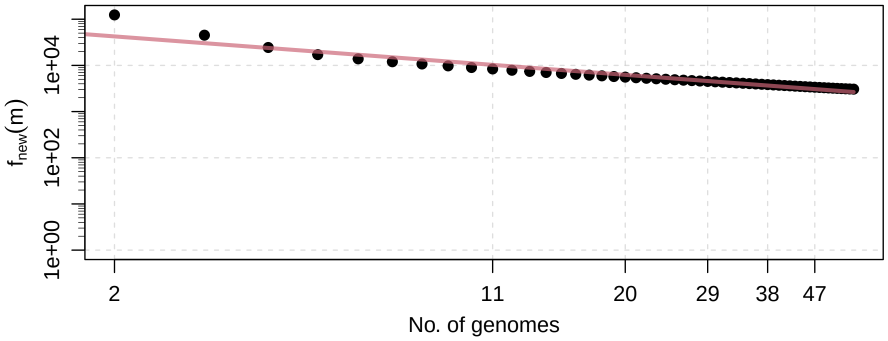

#### S3 Roary

**Figure S21.** Average growth of *Bacillus cereus*.

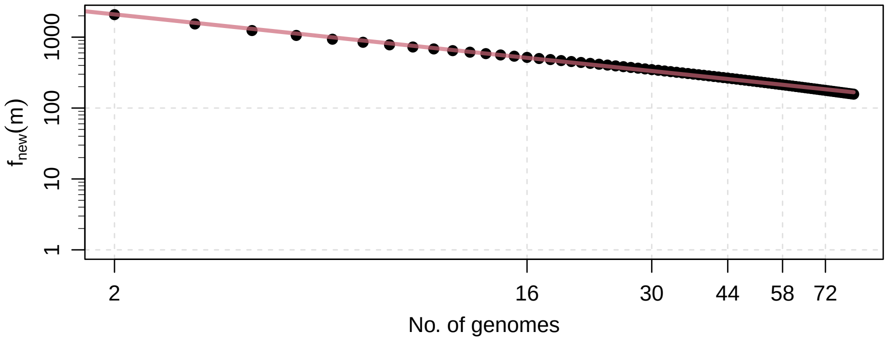


**Figure S22.** Average growth of *Buchnera aphidicola*.

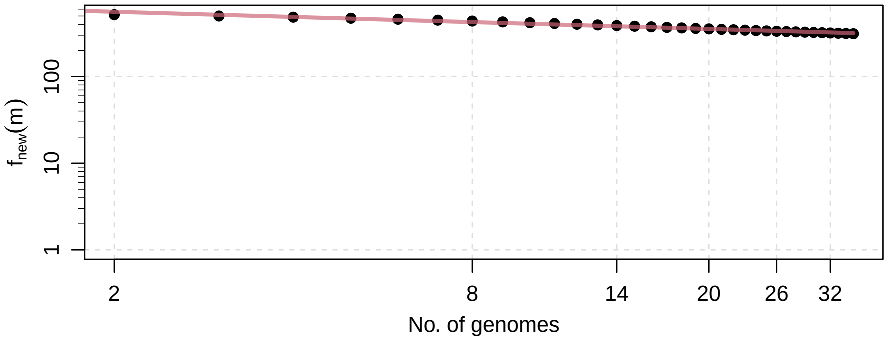


**Figure S23.** Average growth of *Campylobacter jejuni*.

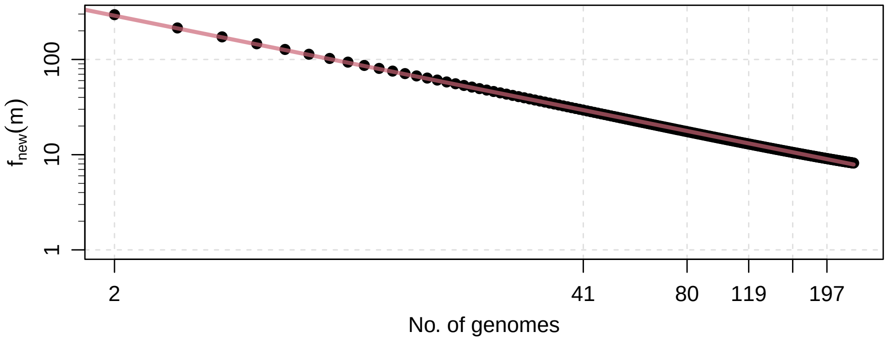

**Figure S24.** Average growth of *Clostridium botulinum*.

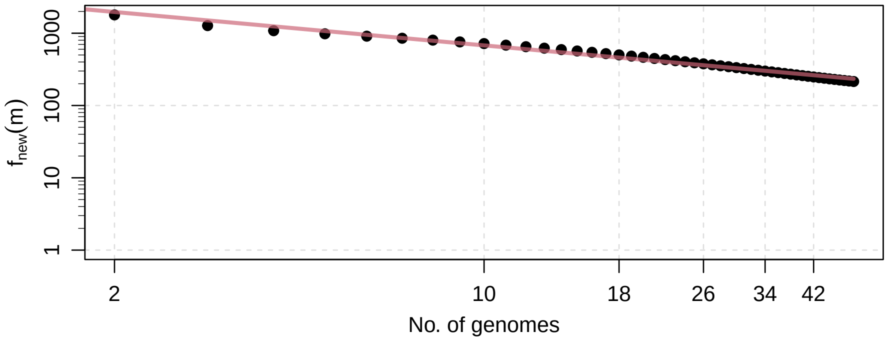


**Figure S25.** Average growth of *Coxiella burnetii*.

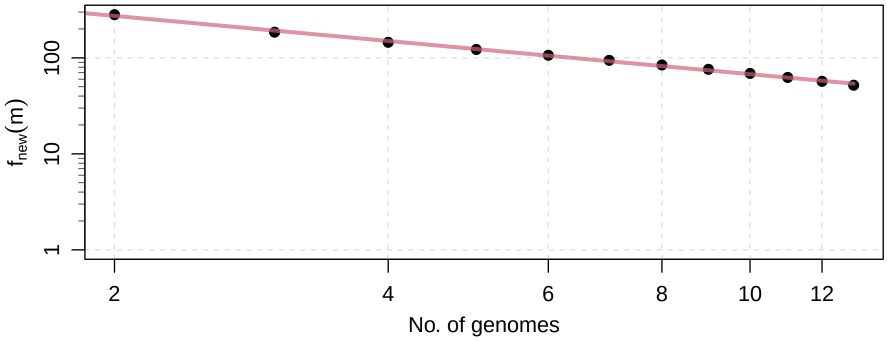


**Figure S26.** Average growth of *Francisella tularensis*.

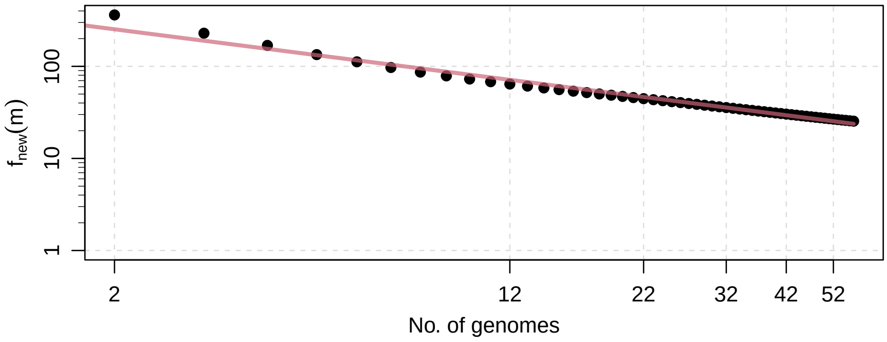

**Figure S27.** Average growth of *Helicobacter pylori*.

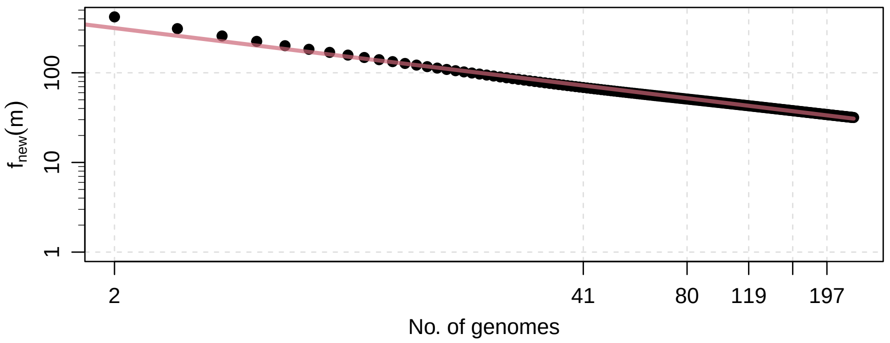


**Figure S28.** Average growth of *Prochlorococcus marinus*.

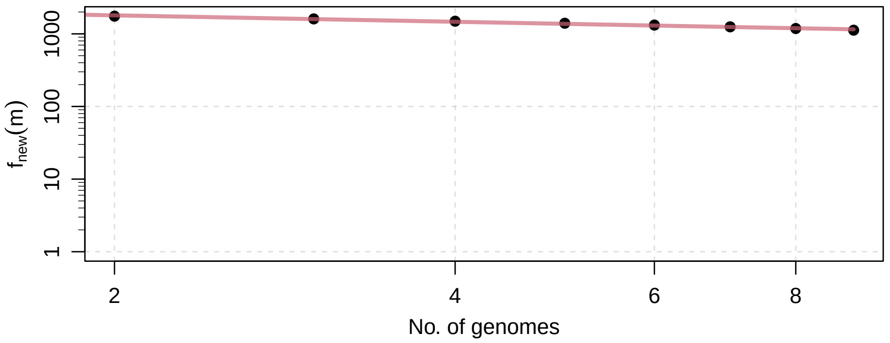


**Figure S29.** Average growth of *Rhodopseudomonas palustris*.

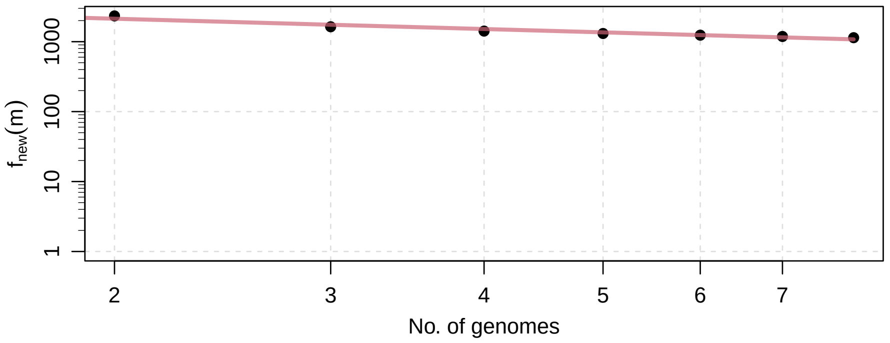

**Figure S30.** Average growth of *Streptococcus pneumoniae*.

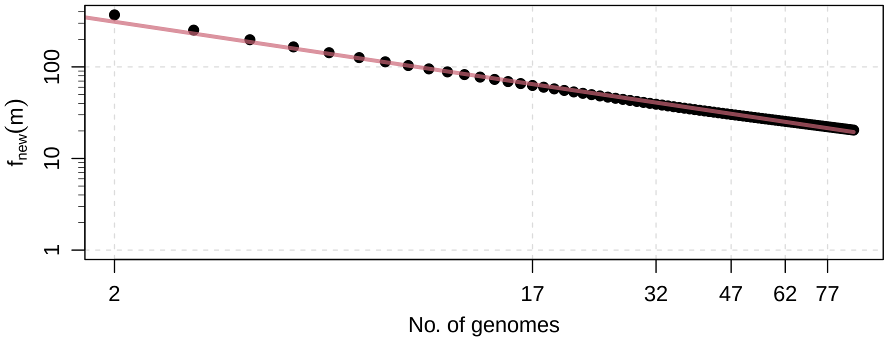


**Figure S31.** Average growth of *Streptococcus pyogenes*.

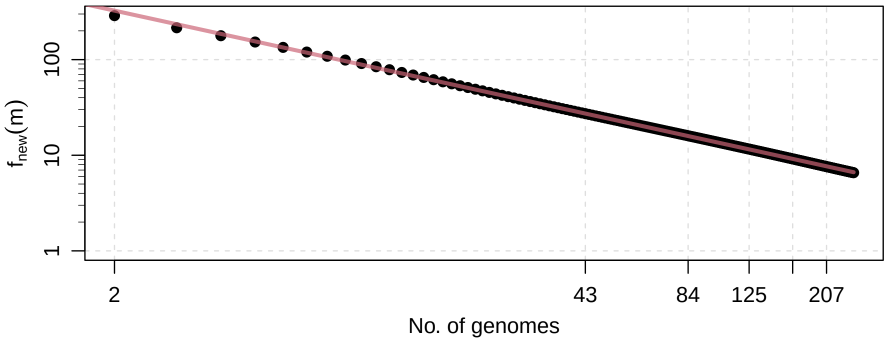


**Figure S32.** Average growth of *Yersinia pestis*.

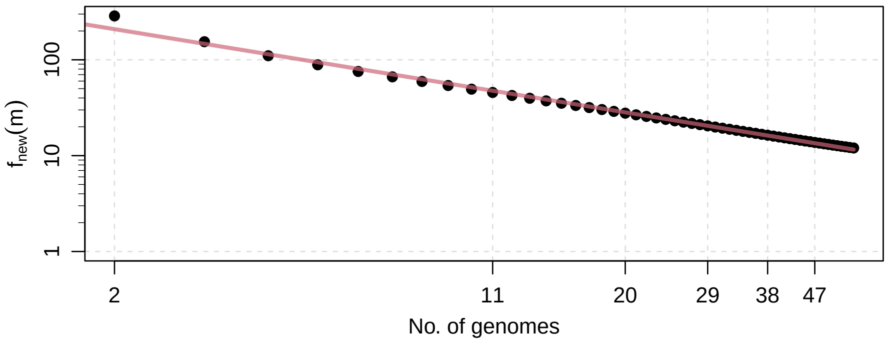

#### S4 Pantools

**Figure S33.** Average growth of *Bacillus cereus*.

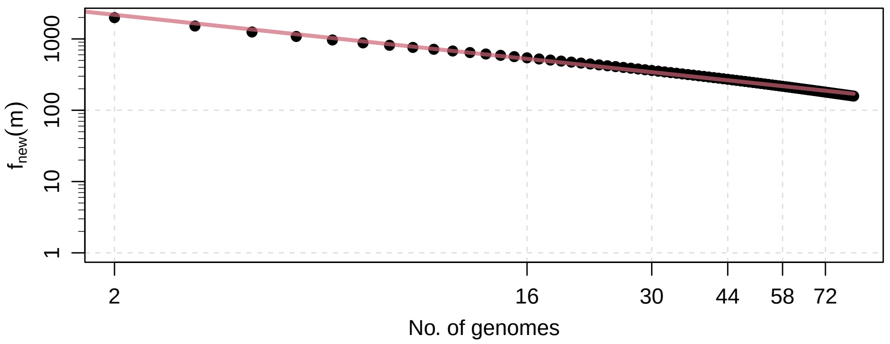


**Figure S34.** Average growth of *Buchnera aphidicola*.

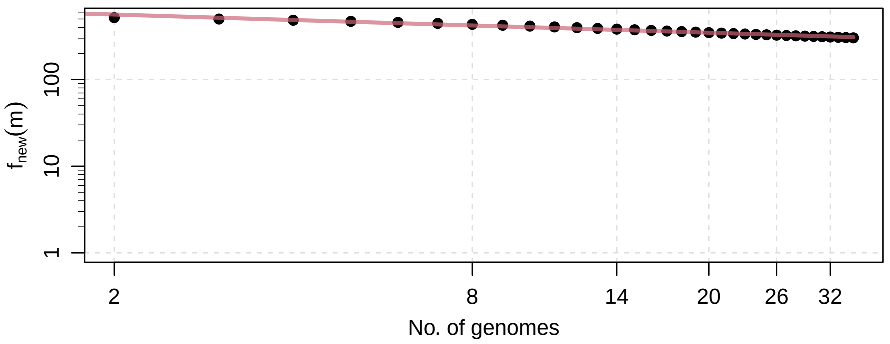


**Figure S35.** Average growth of *Campylobacter jejuni*.

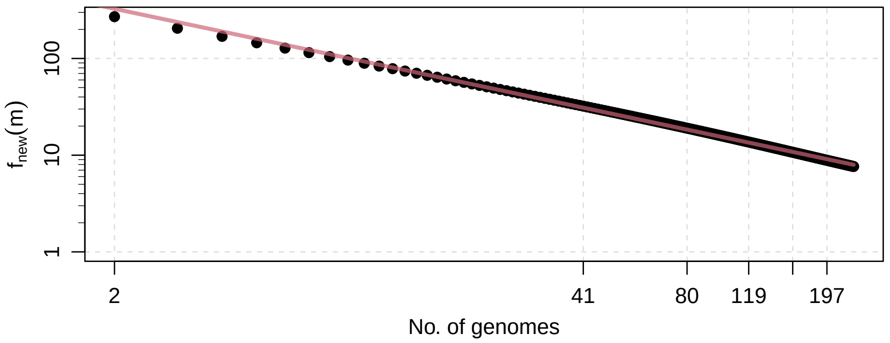

**Figure S36.** Average growth of *Clostridium botulinum*.

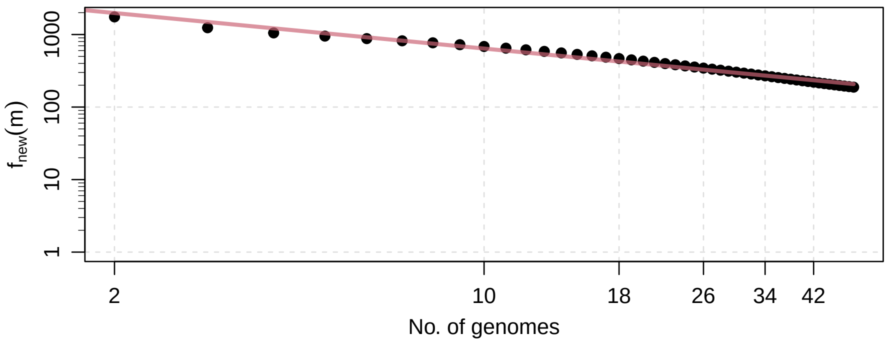


**Figure S37.** Average growth of *Coxiella burnetii*.

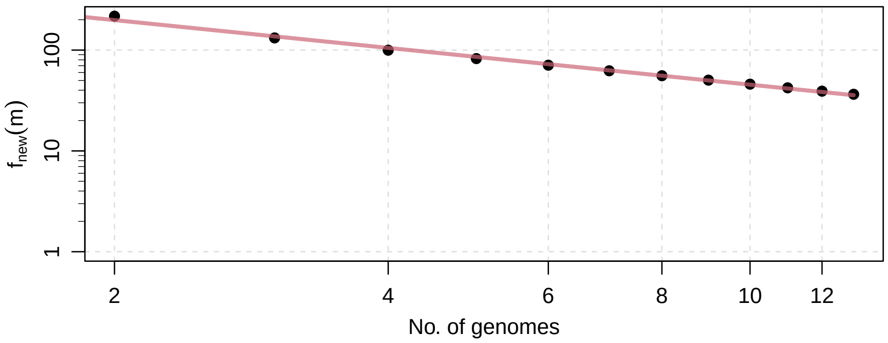


**Figure S38.** Average growth of *Francisella tularensis*.

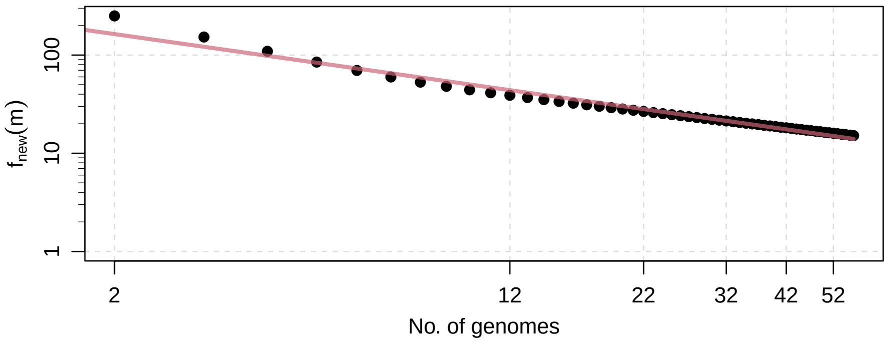

**Figure S39.** Average growth of *Helicobacter pylori*.

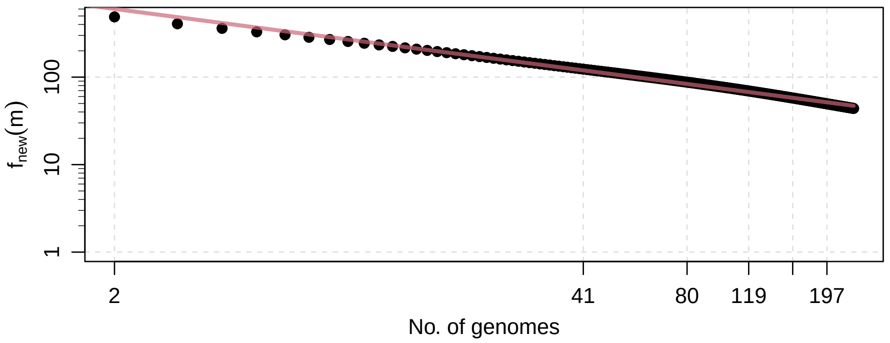


**Figure S40.** Average growth of *Prochlorococcus marinus*.

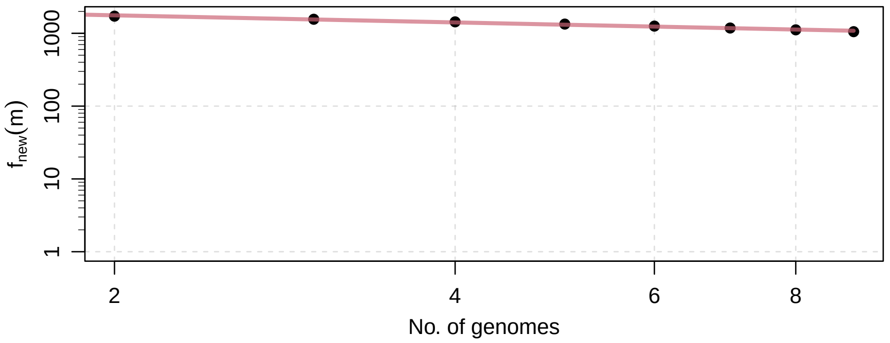


**Figure S41.** Average growth of *Rhodopseudomonas palustris*.

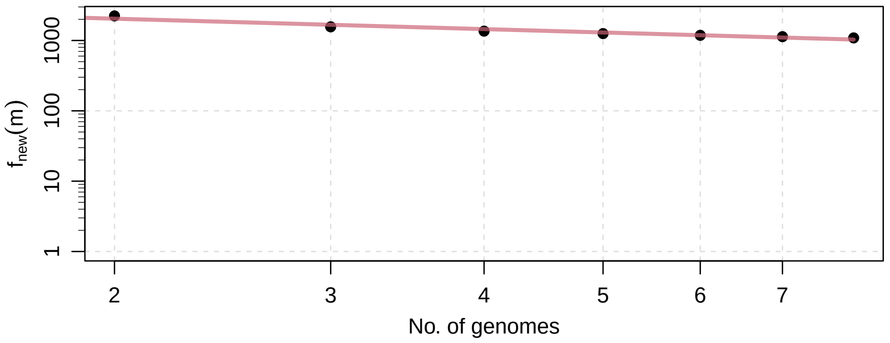

**Figure S42.** Average growth of *Streptococcus pneumoniae*.

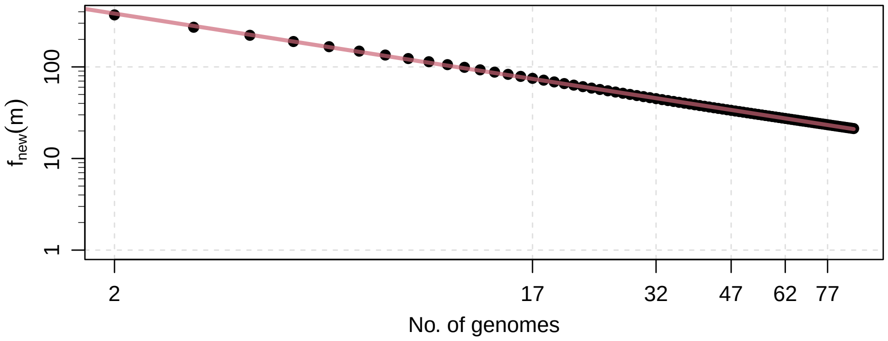


**Figure S43.** Average growth of *Streptococcus pyogenes*.

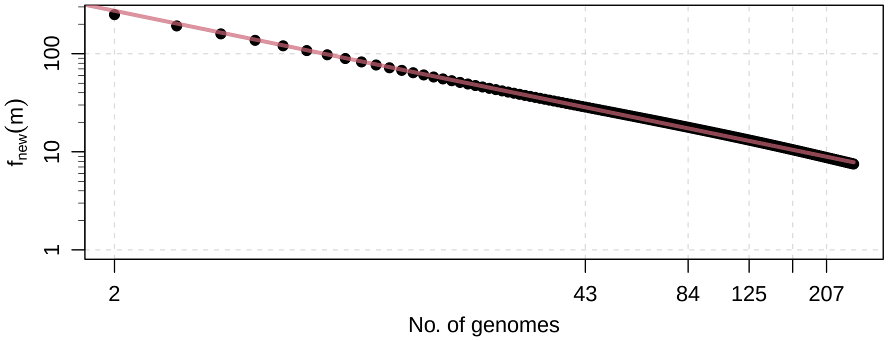


**Figure S44.** Average growth of *Yersinia pestis*.

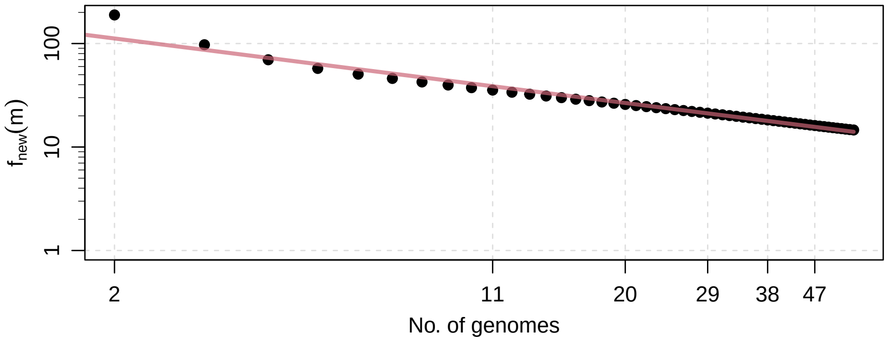

#### S5 BPGA

**Figure S45.** Average growth of *Bacillus cereus*.

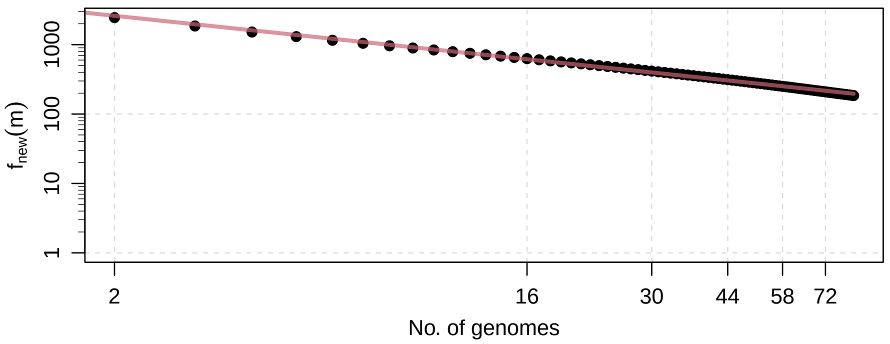


**Figure S46.** Average growth of *Buchnera aphidicola*.

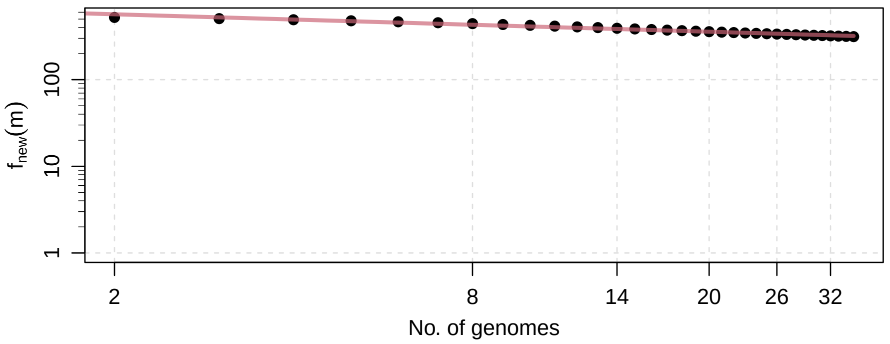


**Figure S47.** Average growth of *Campylobacter jejuni*.

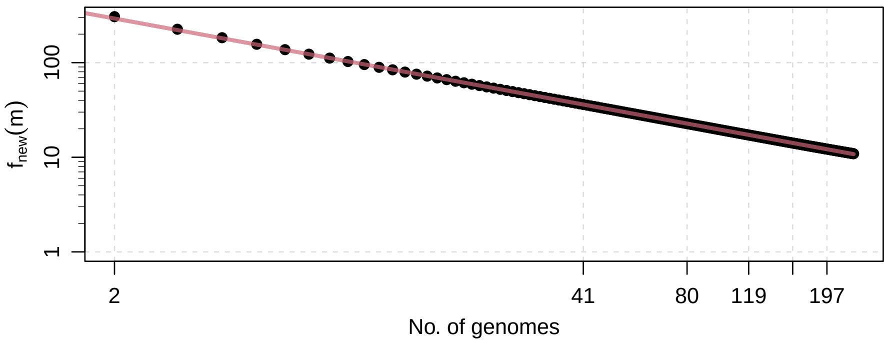

**Figure S48.** Average growth of *Clostridium botulinum*.

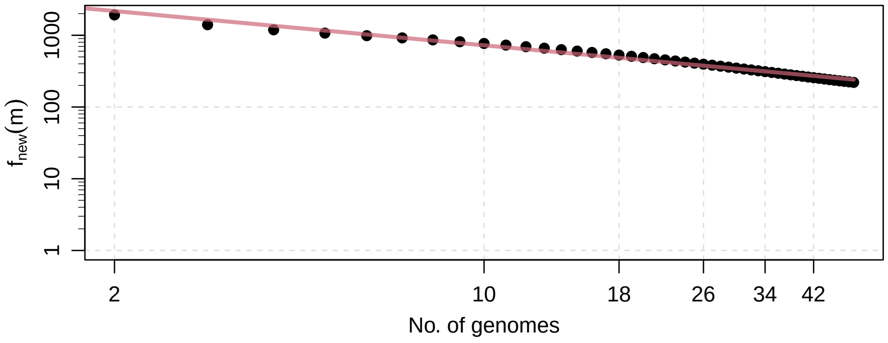


**Figure S49.** Average growth of *Coxiella burnetii*.

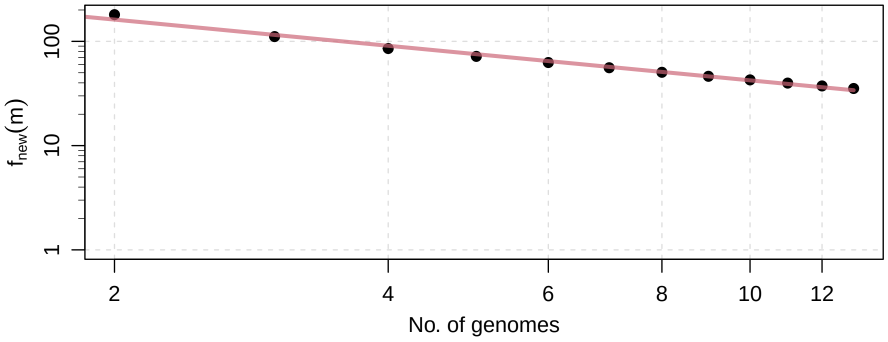


**Figure S50.** Average growth of *Francisella tularensis*.

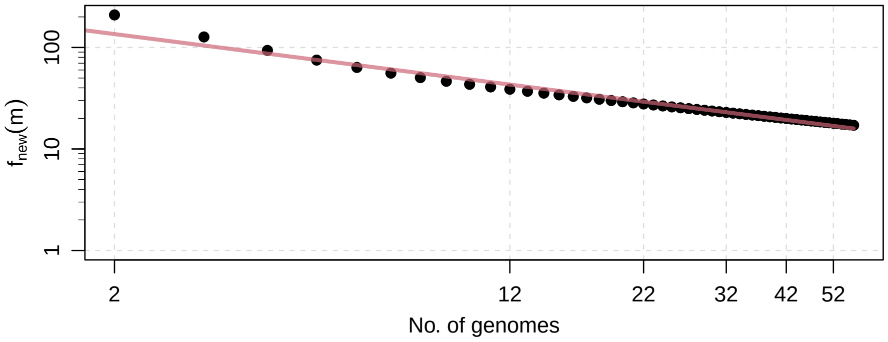

**Figure S51.** Average growth of *Helicobacter pylori*.

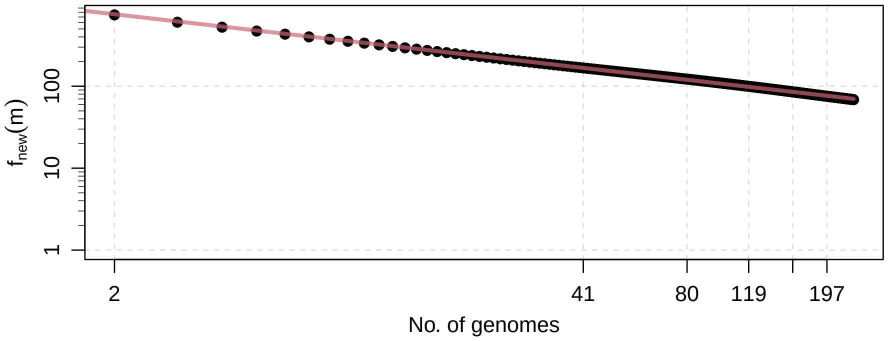


**Figure S52.** Average growth of *Prochlorococcus marinus*.

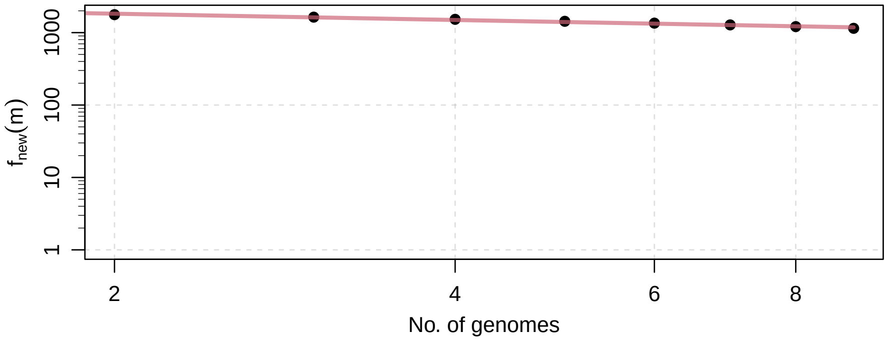


**Figure S53.** Average growth of *Rhodopseudomonas palustris*.

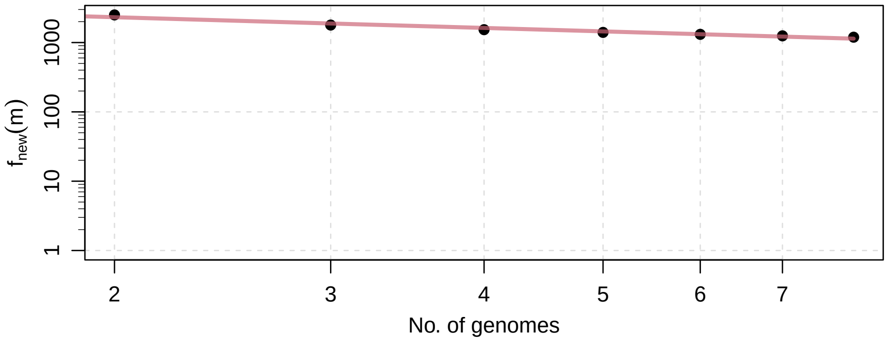

**Figure S54.** Average growth of *Streptococcus pneumoniae*.

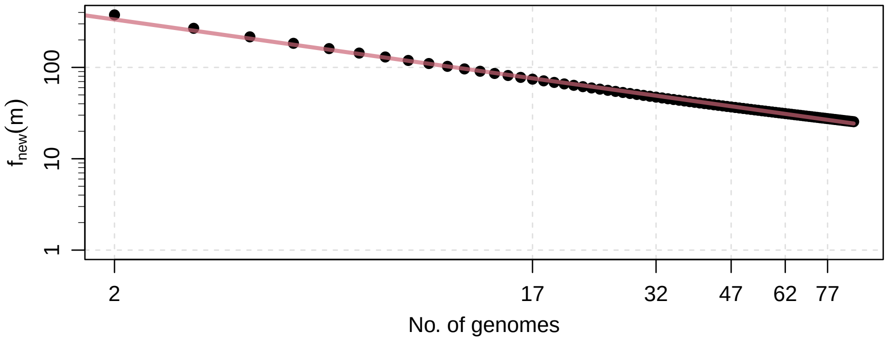


**Figure S55.** Average growth of *Streptococcus pyogenes*.

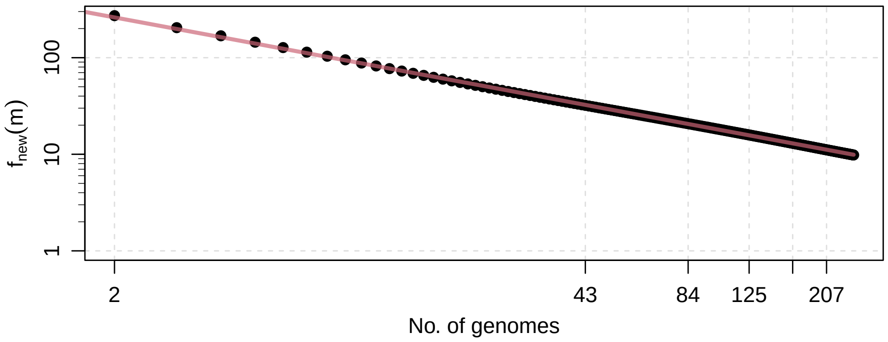


**Figure S56.** Average growth of *Yersinia pestis*.

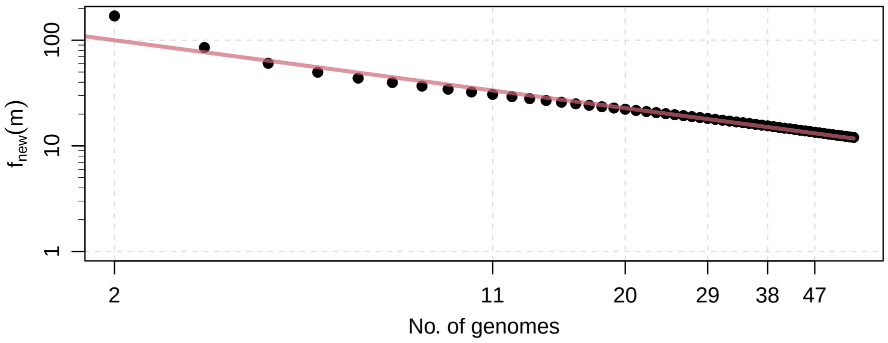

#### S6 Histograms

**Figure S57.** Percentage of items in *Bacillus cereus*.

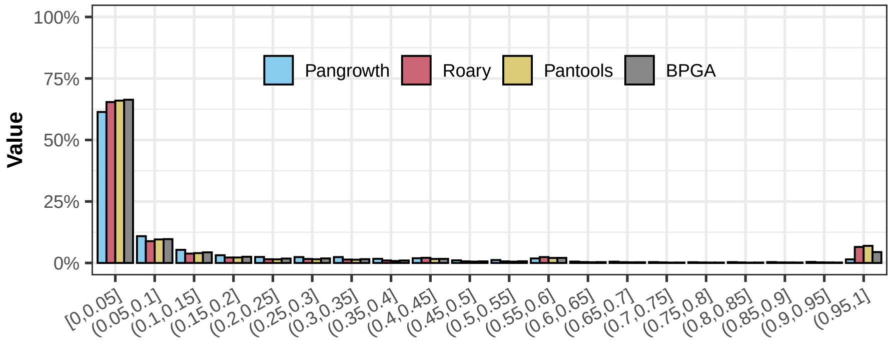


**Figure S58.** Percentage of items in *Buchnera aphidicola*.

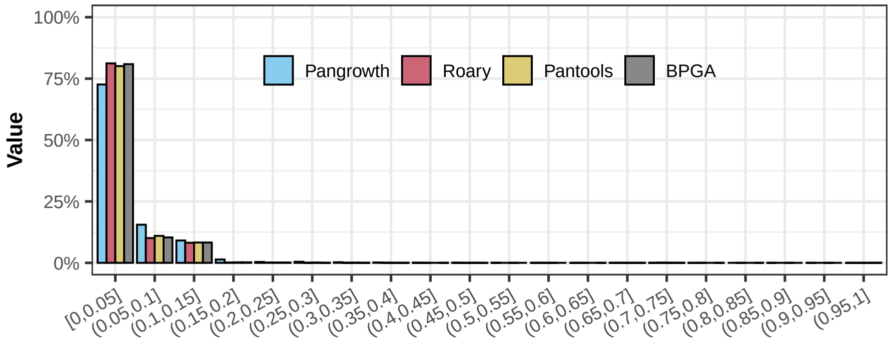


**Figure S59.** Percentage of items in *Campylobacter jejuni*.

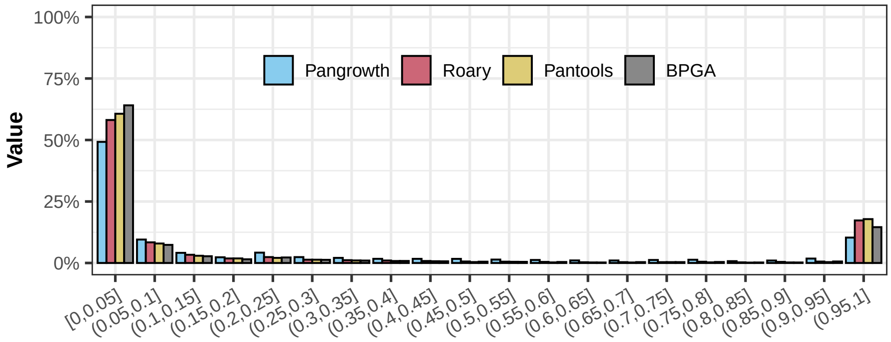

**Figure S60.** Percentage of items in *Clostridium botulinum*.

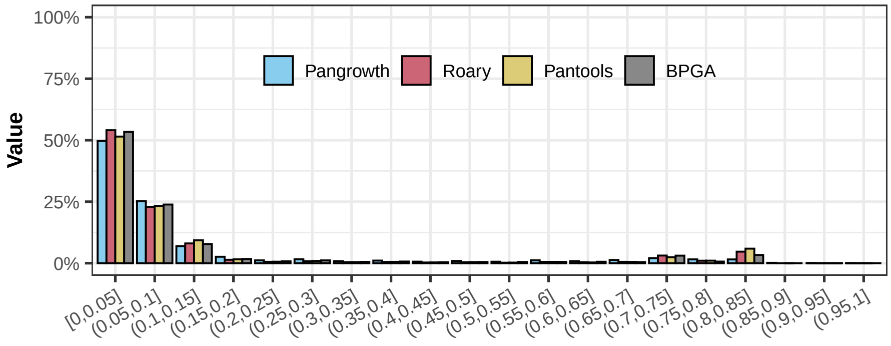


**Figure S61.** Percentage of items in *Coxiella burnetii*.

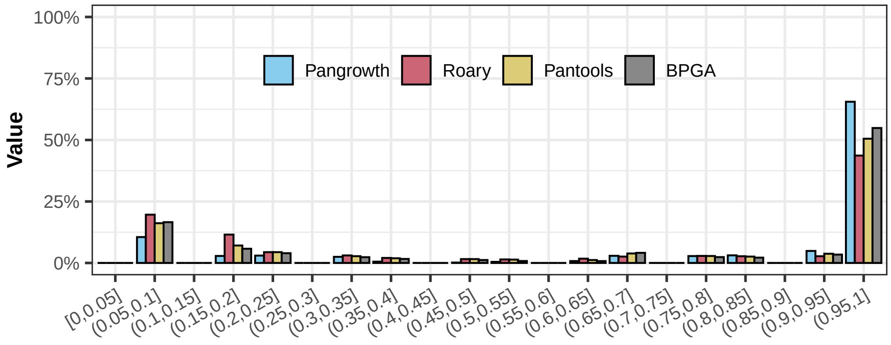


**Figure S62.** Percentage of items in *Francisella tularensis*.

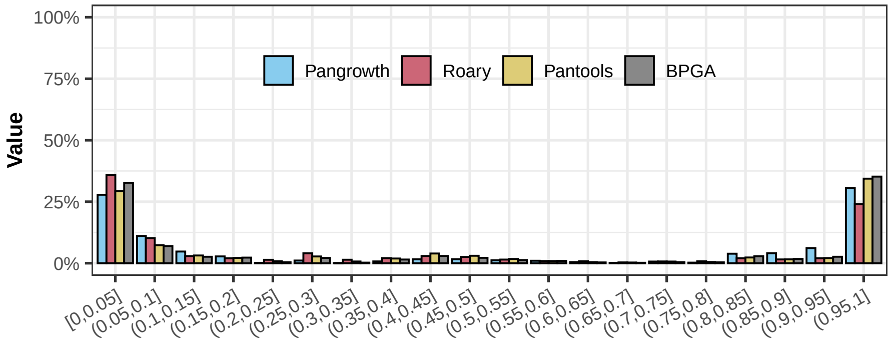

**Figure S63.** Percentage of items in *Helicobacter pylori*.

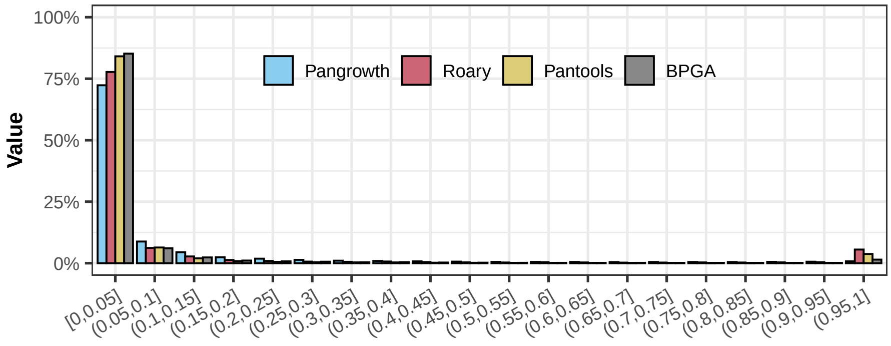


**Figure S64.** Percentage of items in *Prochlorococcus marinus*.

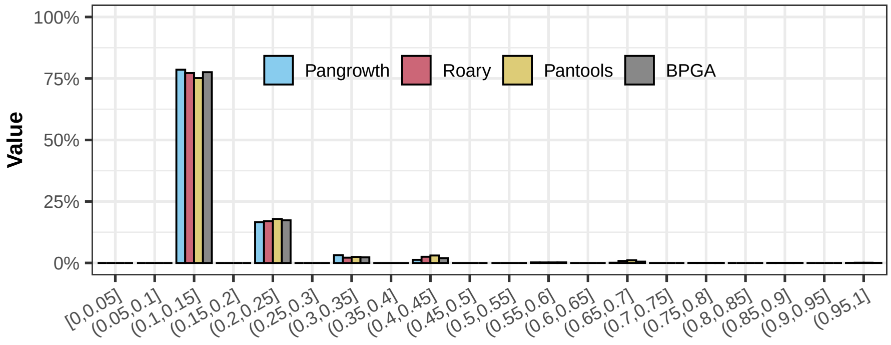


**Figure S65.** Percentage of items in *Rhodopseudomonas palustris*.

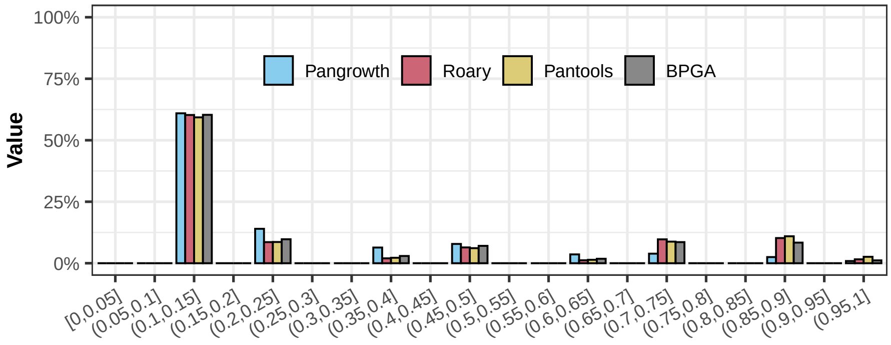

**Figure S66.** Percentage of items in *Streptococcus pneumoniae*.

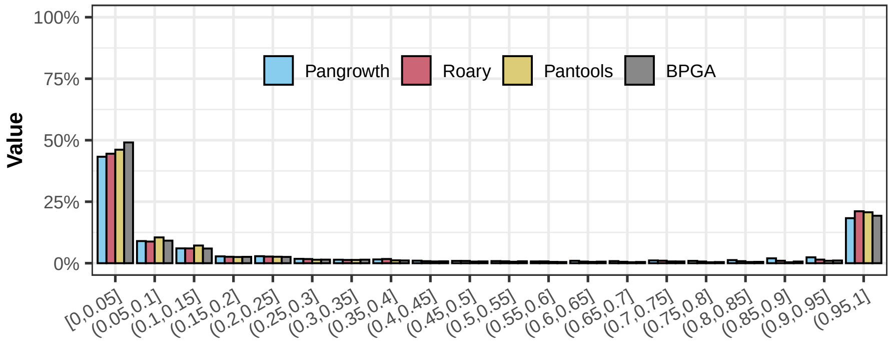


**Figure S67.** Percentage of items in *Streptococcus pyogenes*.

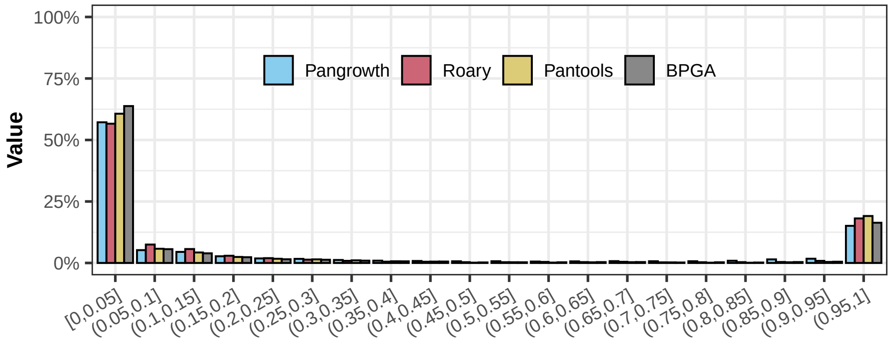


**Figure S68.** Percentage of items in *Yersinia pestis*.

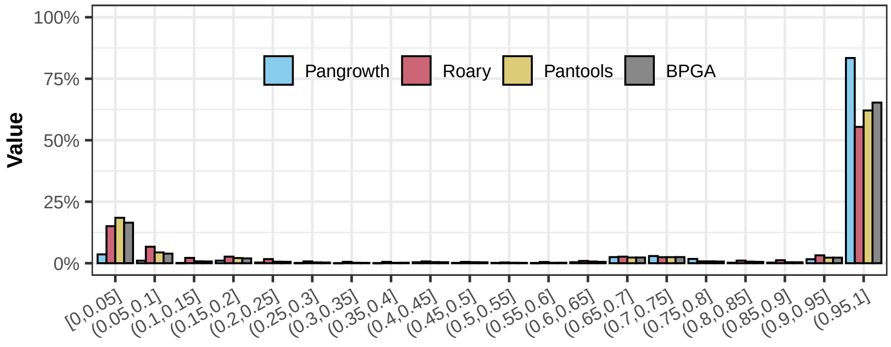
